# Surface-specific film assembly of a Vibrio cholerae adhesin-derived peptide modulated by environmental salts

**DOI:** 10.64898/2026.06.21.733527

**Authors:** Sixin Zhai, Diego R. Jaramillo Pinto, Nicolas L. Mendoza, Adekunle Adewole, Benjamin Heufner, Andrea D. Merg, Tomas P. Corrales, Jing Yan, Roberto C. Andresen Eguiluz

**Affiliations:** Materials and Biomaterials Science and Engineering Graduate Group, University of California Merced, Merced, USA; Department of Chemical and Materials Engineering, University of California, Merced, Merced, USA; Chemistry and Biochemistry Graduate Group, University of California Merced, Merced, USA; Department of Chemistry and Biochemistry, University of California Merced, Merced, USA; Departamento de Física, Universidad Técnica Federico Santa María, Valparaíso, Chile; Department of Molecular, Cellular & Developmental Biology, Yale University, New Haven, USA; Quantitative Biology Institute, Yale University, New Haven, USA; Health Sciences Research Institute, University of California Merced, Merced, USA

**Keywords:** Wet Adhesion, Bacteria Adhesin-inspired Materials, Nano-adhesion, Dynamic Light Scattering, Quartz Crystal Microbalance with Dissipation, Self-assembly Peptide Coating, Interface Science, Surface Science

## Abstract

Underwater adhesion research increasingly draws on bioinspired systems to uncover the molecular mechanisms that enable strong interfacial binding in aqueous environments. The biofilm adhesin Bap1 from *Vibrio cholerae* contains a short peptide motif, SYWFFGWHTK (CP), which exhibits exceptional adhesive performance, surpassing mussel foot protein mfp5 under comparable conditions. Despite its promise, the roles of ionic environments and aggregation behavior in governing CP adhesion remain unclear. In this study, we investigate how ion identity influences CP aggregation, film formation, and interfacial properties. Using dynamic light scattering, we identify the formation of micron-scale assemblies of aggregated molecular clusters (AAMCs), with size distributions modulated by salt type. Quartz crystal microbalance with dissipation and liquid atomic force microscopy reveal that CP film formation is both surface- and ion-dependent. On gold substrates, AAMCs preferentially adsorb and collapse into rigid, smooth nanofilms, consistent with hydrophobic-driven compaction. In contrast, silicate surfaces inhibit such collapse, yielding distinct morphologies and interfacial energetics. These findings demonstrate that surface chemistry and ionic conditions jointly regulate peptide aggregation and adhesion. This work provides mechanistic insight into hydrophobic-rich peptide systems and informs the rational design of next-generation wet adhesives, with broader implications for biomaterials and peptide-based formulations.

**GRAPHICAL ABSTRACT:** 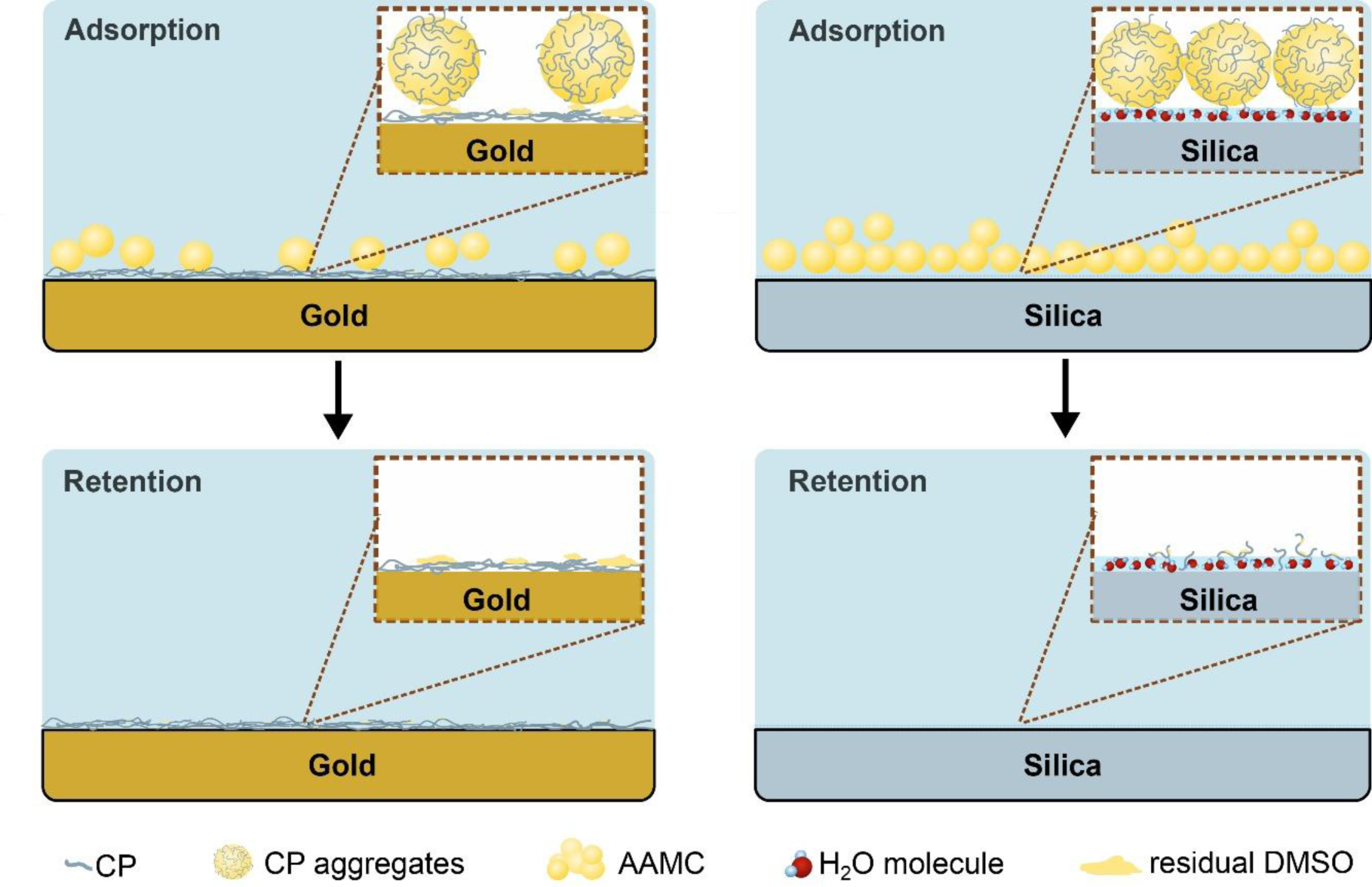

## INTRODUCTION

Underwater adhesive-related research aimed at developing more robust and efficient technologies has focused on elucidating the roles of molecular mechanisms and synergies in biological adhesives across a wide range of species,^[1,2]^ including those derived from bacteria. Bacteria secrete a plethora of extracellular matrix components for strong adhesion to surfaces,^[3,4]^ providing yet another bioinspired library of biochemistries to explore for the development of next-generation wet adhesive technologies. In prior studies of biofilms derived from *Vibrio cholerae*, the causative agent of the pandemic cholerae,^[5]^ we found that the biofilm-specific adhesin Bap1 encodes a peptide sequence that exhibit strong adhesive forces to various surfaces in aqueous environments.^[6,7]^ More specifically, a 10-amino acid domain consisting of SYWFFGWHTK (referred to as CP from now on) (Figure 1) mediated stronger pull-off forces than mfp5, one of the stickiest reported mfps,^[8]^ under similar experimental conditions. Yet, environmental factors and the molecular mechanisms that led to CP’s high adhesive performance require further understanding for the rational implementation onto wet adhesive technologies.

**Figure 1.**
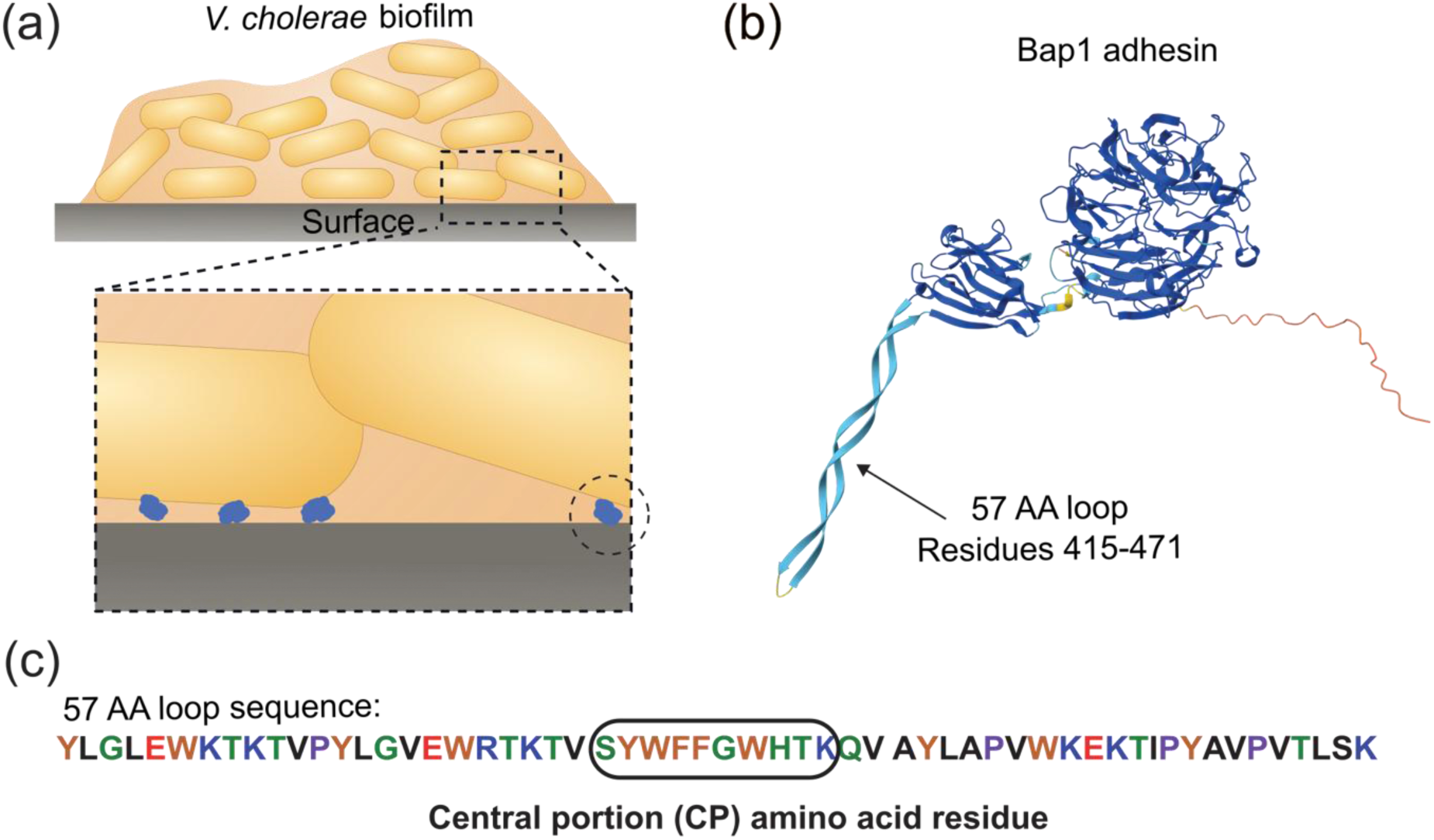
(a) Schematic representing a biofilm of *V*. *cholerae* attached to an abiotic surface showing Bap1, an extracellular matrix adhesin for surface binding. (b) Structure of Bap1 predicted by AlphaFold 3,^[9,10]^ and (c) sequence of the 57 AA loop in Bap1 containing the 10-amino acid central portion (CP) investigated in this study.

Indeed, small molecules^[11,12]^ and peptides,^[13–15]^ such as those derived from Bap1, offer a highly controllable and modular platform for dissecting how multiple weak molecular interactions combine to mediate strong wet adhesion. Effective wet-adhesive peptides typically require a multifunctional design that integrates strong interfacial binding motifs,^[16,17]^ hydrophobic and aromatic residues to facilitate the displacement of interfacial water,^[18]^ and sufficient conformational flexibility to accommodate heterogeneous surface topographies.^[19,20]^ Increasing the complexity of the parameter space, ion-specific (Hofmeister) effects additionally modulate peptide conformation,^[21,22]^ solubility,^[23]^ and interfacial affinity, while increased ionic strength compresses electrical double-layers and influences charge regulation at the interface.^[24,25]^ However, peptide- and protein-based wet adhesive designs are constrained by competing effects: excessively rigid or highly hydrophilic sequences may be unable to effectively disrupt hydration layers or establish contact with the bridging surfaces. Conversely, overly hydrophobic or densely packed sequences can exhibit reduced chain mobility, limiting their ability to reorganize and maximize binding interactions. These trade-offs are further amplified under high ionic strength, where screening can weaken attractive long-range forces while promoting aggregation or collapse.^[26,27]^ This balance motivates the use of protein aggregates,^[28,29]^ dense coacervate phases,^[30–32]^ and related phase-separated systems^[33]^ as efficient adhesives. Such systems enhance local dehydration and intermolecular packing while preserving fluidity and concentrating or excluding specific ions. That level of control allows modulation of interfacial electrostatics and enables efficient surface wetting, spreading, and adaptive binding under ion-rich conditions.

In this work, we explored the impact of ionic nature on the aggregation behavior of CP, with consequences in CP film formation, film morphology, and ultimately, interfacial properties of CP films on different substrates. For this end, we solubilized CP in dimethylsulfoxide (DMSO) and prepared aqueous solutions containing trace amounts of DMSO. Based on dynamic light scattering (DLS), we propose that CP forms micron-sized assemblies of aggregates of molecular clusters, which we call AAMCs, and that the AAMC size distribution in solution is regulated by ionic nature. Additionally, by combining quartz crystal microbalance with dissipation (QCM-D) and liquid atomic force microscopy (AFM), we revealed that CP film formation and interfacial properties induced by AAMCs on gold and silicates were surface- and salt-dependent. From QCM-D experiments, we conclude that AAMCs preferentially bind to gold surfaces, collapsing into rigid nanofilms. From liquid AFM measurements, smooth, dense film morphologies of CP deposited on gold further suggest possible hydrophobic collapse of AAMCs during film formation, in line with our QCM-D observations. These observations contrast with those for AAMCs on silicate substrates, suggesting the absence of hydrophobic collapse and resulting in drastically different interfacial properties compared to those on gold surfaces. These results highlight the dominant role of surface chemistry on CP film formation from AAMCs. Collectively, this work elucidates the interplay among salts, surface chemistry, and CP aggregation, providing insights into the molecular interactions of hydrophobic-rich peptides, with potential applications in wet adhesion technologies, and also contributing to the biophysics of hydrophobic-rich short peptides, relevant to pharmaceutical and drug delivery formulation development.

## RESULTS AND DISCUSSION

### CP formed nano- and micro-scale aggregates in aqueous solutions

The CP peptide in this study contained 50% hydrophobic amino acid moieties, making it challenging to solubilize in aqueous systems (Figure 2a). To overcome similar challenges, biomedical and pharmaceutical research often relies on water-miscible organic solvents. In this regard, DMSO is a commonly used solvent for dissolving water-insoluble peptides and proteins, in part because of its amphiphilic nature, which allows it to associate with both polar and nonpolar moieties.^[34]^

**Figure 2.**
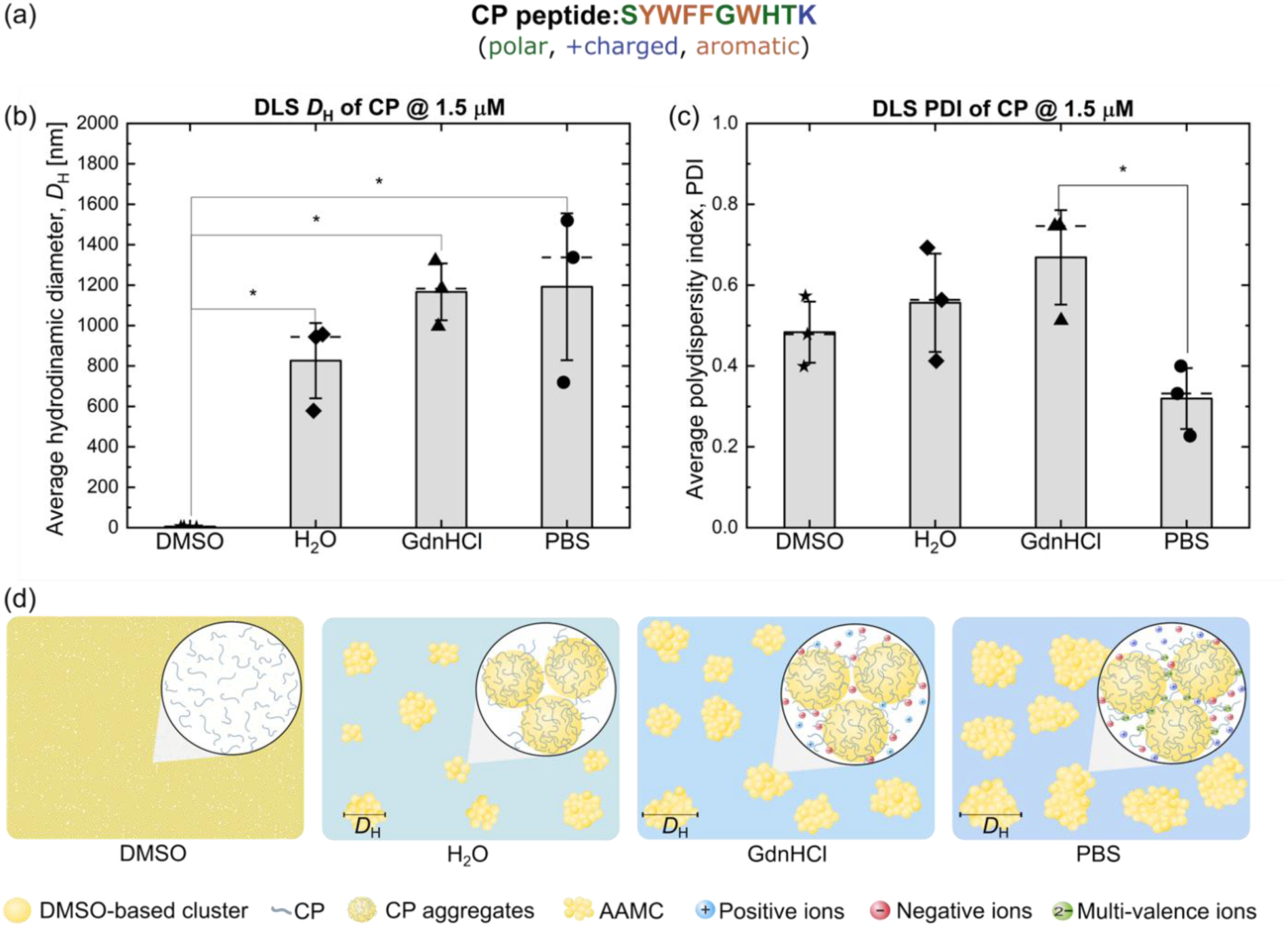
(a) Amino acid sequence of CP indicating moiety properties: polar (green), positively charged (blue), and aromatic (brown). (b) DLS average hydrodynamic diameter, *D*_H_ and (c) polydispersity index (PDI) measured on CP peptide solutions at 1.5 μM in DMSO, H_2_O, GdnHCl, and PBS from 3 independent experiments. Summary tables of data set used for (b) and (c) are displayed in Tables S1 and S3. (d) Schematic depicting peptide aggregates in different solution conditions: pure DMSO, H_2_O, GdnHCl, and PBS. Statistical differences determined by unpaired *t*-test are shown in figures with one asterisk between two conditions if their *p*-values were equal to 0.05 or lower.

To confirm the solubility of CP in DMSO, we performed DLS measurements and extracted the *D*_H_ of CP at 1.5 μM. The mean Z-average *D*_H_ of CP measured in pure DMSO was 5 ± 1 nm, close to the estimated contour length of the peptide (3.5 nm), confirming excellent solubility of CP in DMSO. We then diluted the CP stock solution in water (H_2_O), guanidine hydrochloride (GdnHCl), and phosphate buffer saline (PBS), maintaining a constant concentration of 1.5 μM. A trace amount of DMSO (1 vol.%) was invariably present in each final dilution, which we hypothesize plays a role in AAMC formation. *D*_H_ was approximately 2.5 orders of magnitude higher than in pure DMSO (Figure 2b and Table S1), regardless of the presence or type of salts, as determined by an unpaired *t*-test (Table S2).

To further understand the impact of salt presence and type on CP aggregate size distribution, we report the average polydispersity index (PDI) (Figure 2c). On a scale of 0 to 1, systems having PDI values between 0.1 and 0.7 suggest nearly monodisperse particle size, while a PDI greater than 0.7 indicates a highly dispersed distribution of particle sizes.^[35]^ We consider PDI values as one indicator of CP aggregation behavior in different solutions. The PDI of CP in DMSO was approximately 0.56, with no statistically significant differences and an increasing trend in CP in H_2_O and GdnHCl (Table S3). In PBS, the PDI was the lowest, but only significantly different than the PDI of CP in GdnHCl, as determined by an unpaired *t*-test (Table S4).

As mentioned earlier, CP required initial dissolution in DMSO before further dilution in aqueous solvents. Under hydrated conditions, variations in solvent composition,^[36]^ ionic strength^[37,38]^ and dilution history^[39]^ can drastically affect the assembly morphologies and properties of peptides in bulk solutions, impacting their performance in final products. In general, peptides can undergo self-assembly or aggregation to form nano- to micro-scale structures or large aggregates through various mechanisms. These mechanisms include hydrophobic interactions, electrostatic interactions, hydrogen bonding, cation-π interactions, and aromatic stacking.^[40]^ In our study, we used H_2_O, GdnHCl, and PBS to assess the impact of solvent exchange and salt natures on CP solubility, directly comparing GdnHCl,^[41,42]^ a chaotrope commonly used for denaturing proteins through disrupting hydrogen bonds, and PBS,^[43–45]^ a buffer routinely used in biomedical and pharmaceutical applications as a physiological buffer. Due to its high sodium chloride concentration,^[46,47]^ PBS is considered a kosmotrope in this study.

Based on the DLS measurements, we propose a potential CP aggregation mechanism in aqueous bulk solutions (Figure 2d). *D*_H_ results for CP in DMSO indicated a molecularly dissolved solution. When dissolved in the stock solution, DMSO interacts with both polar and non-polar amino acid moieties of CP, as shown by molecular dynamics simulations in hydrophobic transmembrane helical peptides.^[48]^ Upon dilution in H_2_O, *D*_H_ of CP increased by ∼ 500 times, compared to CP in DMSO. Considering the *D*_H_ measurement of 1 vol.% of DMSO in H_2_O without any peptide (approximately 100 nm)(Figure S1), the large *D*_H_ measured in CP solution in H_2_O containing the same volume of DMSO can be explained as hydrophobic effects driven by hydrophobic amino acid groups in CP (*i.e.*, CP aggregates), to minimize surface exposure to the aqueous environment and free energy.^[49,50]^ Furthermore, CP molecules could also be trapped in DMSO-containing aggregates of molecular clusters. In DMSO-H_2_O binary systems, DMSO induces heterogeneous domains in water due to strong hydrogen bonding, as well as self-interaction. These heterogeneous domains can be aggregates mixed with DMSO-based molecular clusters formed by [(DMSO)_2_], [(DMSO)_3_], [(DMSO)_2_·H_2_O], and [(DMSO)_3_·H_2_O].^[51]^

The micron-sized *D*_H_ of the CP solution in H_2_O could be an AAMC, composed of aggregates of DMSO-based molecular clusters and CP. Similar *D*_H_ values of CP in GdnHCl and PBS suggest that hydrophobic interactions are the major driving force of the assemblies, despite the different salt natures at the working ionic strength (∼160 mM) used in this study. The relatively small PDI measured in PBS, as compared to H_2_O and GdnHCl conditions, suggests the aggregation behavior of DMSO-CP aggregates could potentially be regulated by bridging events between CP via its only cationic moiety, lysine, induced by the divalent cation (HPO_4_^2-^) present in PBS, but more evidence will have to be collected to support the proposed mechanism.

To support the AAMC mechanism, we performed DLS measurements of DMSO-H_2_O, DMSO-GdnHCl, and DMSO-PBS mixtures containing 1 vol.% of DMSO in the absence of the peptides. The average *D*_H_ of all DMSO-based solvent mixtures (Figure S1). The measured *D*_H_ of DMSO-H_2_O mixture, on the nanoscale, is in good agreement with a previous report,^[51]^ but smaller than values measured for CP in water (with the same trace amounts of DMSO). Variations in *D*_H_ measured in DMSO-GdnHCl and DMSO-PBS mixtures compared to the *D*_H_ of DMSO-H_2_O mixture were likely due to changes in the water network or competing hydrogen bonding events between DMSO and H_2_O molecules disrupted or enhanced by GdnHCl or PBS. These measurements revealed the formation of nanoscale heterogeneity in DMSO-based solvents prior to peptide addition. Although these nanodomains were substantially smaller than the micron-sized AAMCs, they may modulate peptide solvation and local intermolecular interaction that likely governs the nucleation of CP aggregate assemblies (AAMC). Overall, we anticipated aromatic stacking or cation-π interactions among aromatic AA moieties, including FF, which are known to drive beta-amyloid aggregation.^[52]^

### CP films formed faster, denser, and more stable on gold than on silica substrates

A key property of an effective adhesive is that its molecular constituents have a high affinity for the surface to which it bonds. Using QCM-D, we compared the impact of substrate chemistry on CP film formation across different aqueous environments with varying salt nature and concentration (Figure 3).

**Figure 3.**
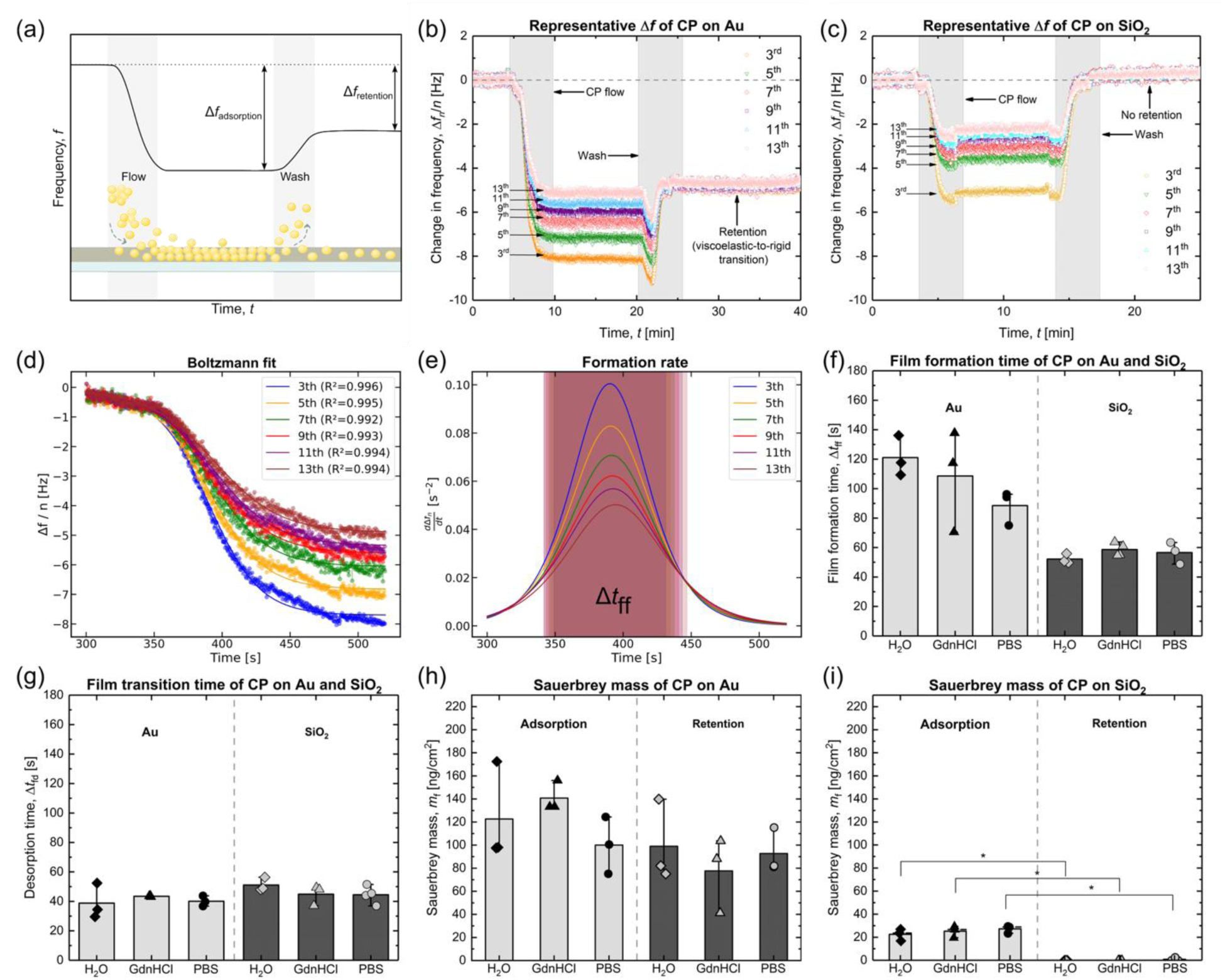
(a) Schematic of QCM-D changes in frequency for a typical CP experiment. (b) and (c) Plots of representative changes in frequency Δ*f_n_*/*n* over time, where *n* represent the overtone (from the 3^rd^ to the 13^th^) measured by QCM-D for CP in PBS forming films on gold and silica substrates, respectively. Representative Δ*f* plots of CP on gold and silica in H2O and GdnHCl are shown in Figure S2. (d) Representative sigmoidal fits of Δ*f_n_*/*n* versus time for CP in PBS on Au. *R*^2^ values for the fits of each overtone are reported. (e) Derivative of the sigmoidal fits as a function of time to obtain the film formation times. (f) and (g) Average film formation (CP flow), and transition (wash) times, respectively. For (f) and (g) each data point represents an average calculated following the steps on (d,e). (h) and (i) Sauerbrey mass *m*_f_ calculated from the steady states pre-wash (adsorption) and post-wash (retention) on gold and silica, respectively. Differences between *m*_f_ of CP in different solvents and on substrates were evaluated using two-way ANOVA tests. Differences between *m*_f_ of CP adsorption and retention were evaluated using an unpaired *t*-test. Statistical differences between two conditions are displayed with one asterisk above them on the plot if their *p*-values were 0.05 or lower.

Substrate nature (*i.e.*, gold versus silica) had a significant impact on the frequency changes when flowing CP peptide solutions, regardless of solvent conditions (Figure 3b and 3c). For both types of substrates, after deposition (adsorption step, pre-wash), each overtone showed different frequency changes (*i.e*., a spread between the overtones). This indicates two characteristics of the films: they were anisotropic in the direction normal to the sensor, and they were not completely rigid/elastic. For gold, film thinning was evident after the wash, with all overtones collapsing onto a master curve, indicating a transition from a soft/viscoelastic heterogeneous layer to a rigid, more homogeneous one. This is also evident on the dissipation channel (Figure S3). On gold, the value to which all overtones collapsed was close to the value of the 13^th^ overtone. The fact that all overtones collapsed to the value of the 13th overtone suggests that the lower part of the film remained strongly bound to the surface, while the upper layers were either removed or underwent structural collapse. That is, as in QCM-D, different overtones “probe” films at different depths and widths. Lower overtones (*e.g.*, 3^rd^ and 5^th^) probe deeper and over larger areas of the coupled film, as those overtones have the highest penetration depth and width. Higher overtones (*e.g.*, 11^th^ and 13^th^) probe film properties near the sensor surface. Interestingly, on silica, an initial viscoelastic CP film formed, but was easily removed after the wash step, with no measurable film mass.

Next, to better understand peptide-substrate interactions, we studied film-forming kinetics. For this end, changes in frequency measured by QCM-D for overtones 3^rd^ to 13^th^ during film formation (CP flow) and desorption (wash) were fitted to a sigmoidal Boltzmann function (Figure 3d and Figures S4 to S7). Once fitted, we calculated formation rates from the time derivatives of each Boltzmann fit (see the Materials and Methods section for a detailed explanation), revealing a clear overtone dependence (Figure 3e). We averaged values extracted for each overtone and used their means to inform on film formation and desorption times (Figures 3f and 3g). Film formation time between substrates differed, as supported by two-way ANOVA, being ∼5 times longer on gold than on silica, independent of solvent. On the other hand, desorption times (*i.e.,* the time it takes to remove loosely bound CP aggregates) were similar on both substrates and across solvents.

To further obtain information on CP films formed on gold and silica substrates, the adsorbed and retained mass density, *m*_f_, was quantified (Figures 3h and 3i). *m*_f_ of CP on gold was similar across H_2_O, GdnHCl, and PBS, for both adsorption and retention. On silica, this was also true; however, no CP *m*_f_ after the wash (*i.e.*, retention) was measured. Differences between *m*_f_ of CP adsorption and retention were evaluated via a paired *t*-test (Tables S5 and S6). Differences between *m*_f_ of CP in different solvents and on substrates were evaluated using two-way ANOVA tests (Tables S7 and S8).

On gold, CP films before wash had an *m*_f_ about five times that of on silica. After the wash, the difference in CP film retention strength for both substrates became evident, as most of the film was retained on gold for all conditions, but not on silica (adsorption and retention steps were significantly different across solvent types on silica). Gold, as a more hydrophobic substrate than silica, forms weaker hydration barriers for CP. CP, composed of 50% hydrophobic amino acids, was likely collapsing from its aggregated state onto the gold surface, forming multiple contact points that rearranged to a lower-energy state and assembled into a thin, densely packed film that favored interactions with the gold surface. Before washing, energy-dissipative aggregates in the form of AAMCs were loosely bound to the thin, rigid CP layer. These aggregates were washed away while the thin peptide film remained bound to the surface. As discussed above, the chaotropic nature of GdnHCl would be anticipated to impact, to some extent, hydrophobic interactions, compared to H_2_O or PBS. However, at the working concentration of GdnHCl, disruptions in the hydrophobic peptide-gold interactions were not observed, at least from QCM-D insights. The mass density of the retained films was consistent with the theoretical monolayer mass density of CP, as shown in the supplemental information (S1. Estimation of CP monolayer mass density).

On silica, strong hydration barriers are present at the sensor surface,^[53]^ preventing direct interaction of AAMCs with silica. This inhibited the hydrophobic collapse of AAMCs onto the surface, thereby preventing the formation of a CP film. Our interpretation (Figure 4) is that no continuous or homogeneous peptide film formed, with AAMCs deposited on top of the hydration layers during adsorption, which were completely removed after washing. However, a few CP molecules that penetrated the hydration layer remained bound (albeit undetectable by QCM-D) and have the capabilities to mediate strong pull-off forces, as previously reported.^[6]^

**Figure 4.**
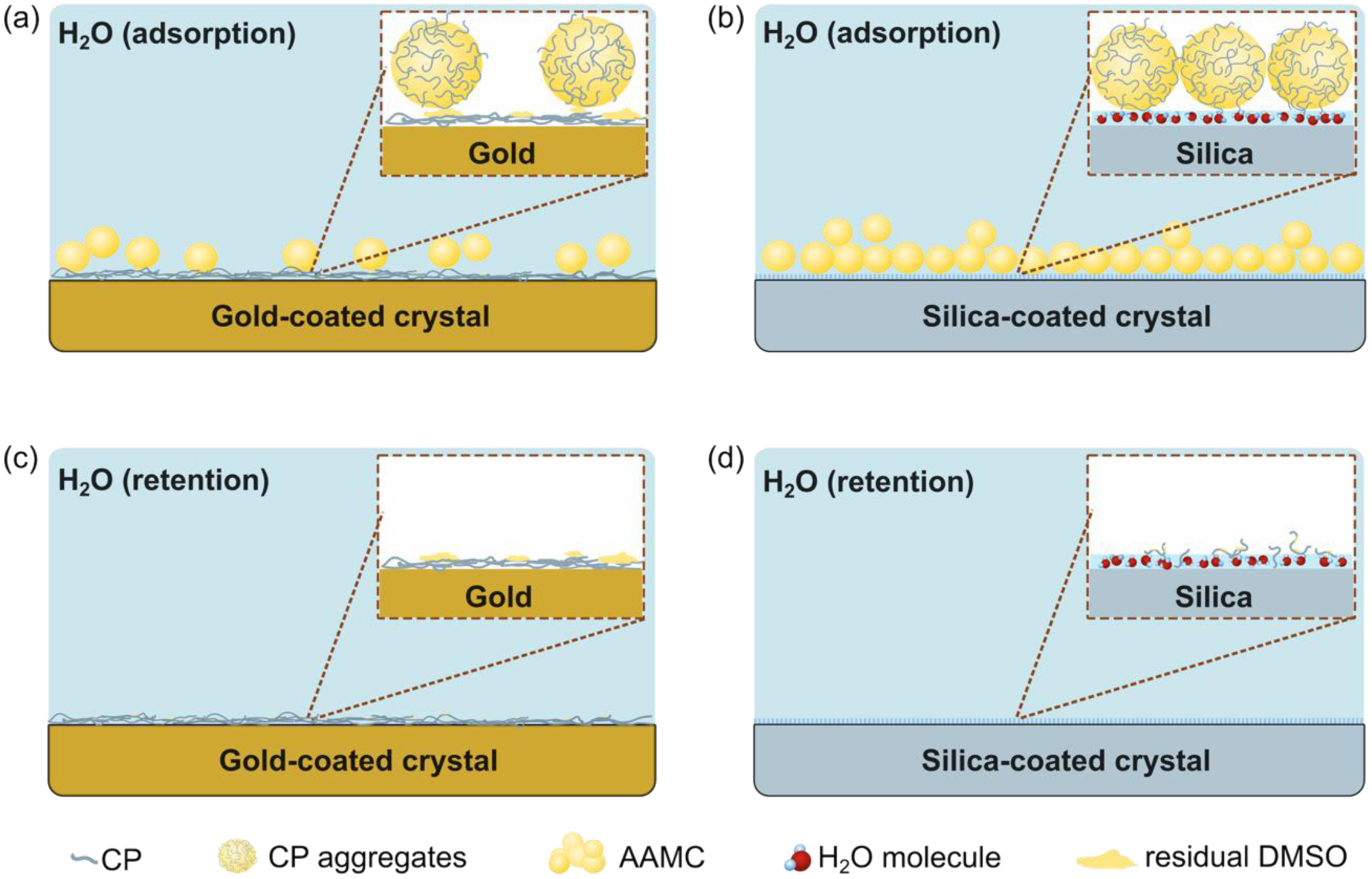
Schematic diagram of stabilized adsorption and retention phases of CP-exposed QCM-D crystals in H_2_O, showing (a) AAMC deposition on CP films formed on gold-coated crystals and (b) AAMC on hydration layer formed on silica-coated crystals during adsorption. After washing, AAMCs are removed (c) leaving a thin, rigid CP film on the gold surface

Overall, CP films on gold were ∼ 5 times denser than on silica, and the formation time of CP films on gold was ∼ 2 times slower than for silica. This indicates that CP film formation occurred at a slower rate on gold than on silica. However, desorption times for gold and silica were similar, suggesting that loosely bound aggregate structures that did not collapse (for gold or silica) were easily removed during the washing steps. These results reveal that surface chemistry plays a crucial role in mediating the molecular assembly of CP films, as has been reported for other protein and peptide^[54]^ systems rich in hydrophobic content, such as Mfps.^[55–57]^

### Retained CP films on gold were dense, smooth, and rigid, with solvent-influenced surface morphologies

From DLS and QCM-D measurements, we gained insights into the bulk aggregation of CP in different aqueous environments and how this affected film formation/desorption kinetics and amounts, depending on surface chemistry. To further confirm our understanding of CP film morphology and gain information on their nanomechanical/surface properties, we used AFM in liquid environments. All measurements presented are representative of at least 3 independent samples.

On gold, CP in water was the most challenging condition to image. The height channel (Figure 5a), collected by QI imaging from force-distance curves of interactions between the AFM probe and the sample, shows a relatively smooth surface with low heterogeneity. The difficulty in resolving high-detail surface morphology arose from complex tip-sample interactions, evidenced by long-range interactions during approach and retract force-distance (FD) curves (Figure 5d). The presented FD curve shows an abrupt jump into an attractive regime ∼100 nm before contacting a rigid surface, presumably a CP film. While in the attractive regime, the FD curve shows two instantaneous jumps ∼ 100 pN, suggesting a potential layer penetration as the probe indents. We hypothesize that residual DMSO might be binding to hydrophobic domains of CP, creating a nanostructured layer, resting on top of a rigid CP film, in agreement with QCM-D observations.

**Figure 5.**
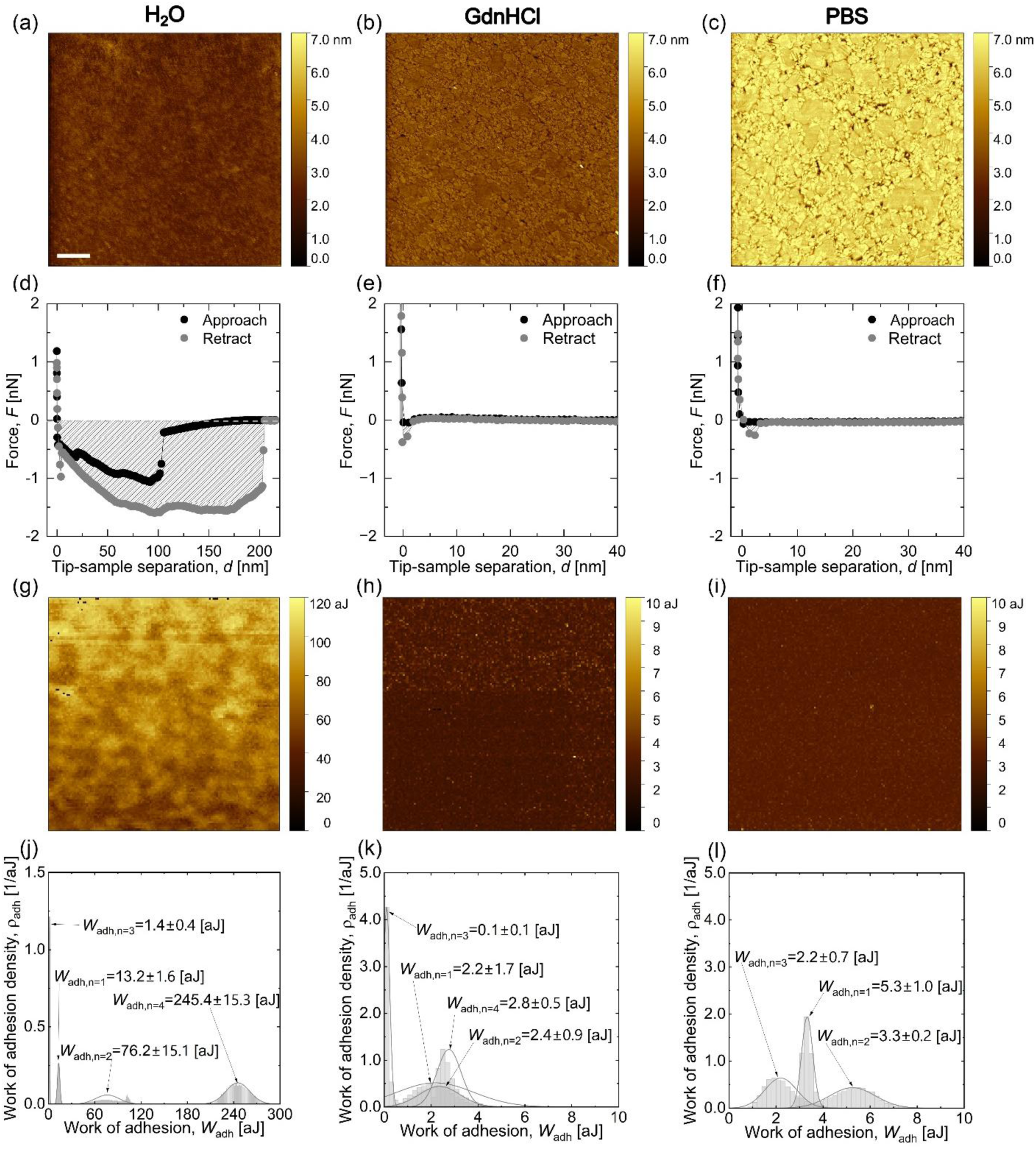
Representative AFM measurements of CP films on mica-templated gold in different aqueous environments. Height maps of CP films in (a) H_2_O, (b) GdnHCl, and (c) PBS, respectively. Representative force-runs in (d) H_2_O, (e) GdnHCl, and (f) PBS, respectively. Work of adhesion (*W*_adh_) maps of CP films in (g) H_2_O, (h) GdnHCl, and (i) PBS, respectively. Note that the color scale of *W*_adh_ maps were adjusted to optimized visualization of local differences in interfacial energies measured. (j), (k), (l) *W*_adh_ distribution histograms of CP films extracted from the *W*_adh_ maps from all measurements. Scale bar in (a) is 500 nm and applies to all AFM maps shown.

The DMSO nanolayer would naturally create two extra interfaces: *i*) between the probe (immersed in water) and DMSO, and ii) between DMSO and CP film, explaining the consecutive “jumps” of the FD curve for both approach and retract. Further evidence supporting this hypothesis is provided in the supplemental information (Figure S8), which shows two populations of jump-out forces measured as the tip retracted from the CP-DMSO film in the vertical *z* direction. To further explore the idea of residual DMSO binding to CP, we show the work of adhesion (*W*_adh_) map (Figure 5g), which quantifies the negative area of the retract FD curve (filled area in Figure 5d). *W*_adh_ includes contributions from the adhesive and viscoelastic behavior of the films, as well as interactions with the tip (if any). The heterogeneous patches revealed in the *W*_adh_ maps (Figure 5g), but absent in the height maps, suggest that DMSO forms interconnected domains on top of CP. We additionally show histograms of *W*_adh_ obtained from all independent CP measurements in H_2_O on gold (Figure 5j). The condition of CP in H_2_O on gold was, out of all reported conditions, the one displaying the highest magnitudes and the widest ranges of *W*_adh_. Typical height maps of CP in GdnHCl and PBS on gold (Figures 5b and 5c) reveal that the surface morphology clearly resembles that of bare mica-templated gold (Figure S9). This agrees with our previous interpretation of a compact, dense CP monolayer adsorbed to gold substrates, as inferred from QCM-D. In both cases, FD curves show smooth contact with short-range interactions between the AFM probe and the CP films, which were too stiff to be deformed under the applied load (in excellent agreement with QCM-D interpretations). *W*_adh_ distributions were lower for GdnHCl and PBS than for H_2_O, as evidenced by the *W*_adh_ maps (Figures 5h and 5i) and by more consistent normal distributions (Figures 5k and 5l). No evidence of DMSO nanolayers was observed, contrasting with CP in H_2_O and, to our interpretation, leading to lower *W*_adh_ values. As quantified by QCM-D, the mass density differences between CP films deposited in different solvents were not significant. However, subtle differences in the CP film-forming mechanism, attributed to solvent composition, may exist. It is possible that when forming films in water, with an ionic strength of ∼ 0, AAMCs from bulk solution were able to collapse onto the gold surfaces due to the hydrophobic effect, as previously discussed, in a preferred conformation that allowed DMSO molecules to be kept attached to hydrophobic (aromatic) domains of CP. For GdnHCl and PBS, on the other hand, the salt ions in solution might have interfered with this process via cation-*π* interactions, forcing DMSO molecules to detach from CP once the AAMCs collapsed to form the monolayer, resulting in a DMSO-free film. This might be the reason for the slightly higher retained mass density of the CP films in H_2_O compared to the other solvents (Figure 3h), although that difference was not enough to be considered statistically significant by the applied unpaired *t*-test. In other words, even if the mass density is similar for CP across solvents on gold, the ions in solution prevented DMSO from binding to CP hydrophobic domains, disrupting the formation of DMSO nanodomains on the CP films in GdnHCl and PBS. Next, we discuss the AFM results of CP deposited on mica.

### Retained CP films on mica presented morphological differences across solvents

Mica and silica have been extensively used as analogous model surfaces, as they are both hydrophilic silicate-based materials bearing negative charges in water, that form similarly strong hydration layers^[58]^. Additionally, using mica as a substrate minimizes roughness artifacts that could affect CP adsorption.^[59]^ Note that we did not use mica in QCM-D since functionalizing QCM-D sensors with mica is possible but challenging, often resulting in a compromise in mass sensitivity and/or overtone accessibility.^[60]^

As surface roughness is known to be a contributor regulating peptide adsorption to surfaces, we used mica-templated gold (measured surface roughness: ∼ 0.6 nm) as a more hydrophobic model substrate for liquid AFM compared to mica (measured surface roughness: ∼ 0.3 nm) to mitigate the influence from surface height. Across conditions, the morphology of CP films on mica was significantly different from that on gold, confirming the importance of substrate chemistry, as observed via QCM-D and supported by root mean square (RMS) roughness measurements (Figure S10 and Table S9). We observed that CP formed dense and almost continuous films composed of CP aggregates (∼50 nm in diameter) that retained globular shapes in H_2_O and GdnHCl, respectively (Figures 6a and 6b and controls without CP in Figure S11). Connecting with DLS measurements, the AAMC structures seemed to have dissociated into their smaller constituents (CP aggregates). The morphological difference compared to gold may be due to mica forming stronger hydration layers, as detailed in the QCM-D section. In contrast, CP in PBS showed micron-sized structures that discretely decorated the mica surface. We interpret this finding as AAMCs that did not dissociate into CP aggregates, stabilized by divalent ions present in PBS, and that were not washed away, in contrast to QCM-D silica observations. To reconcile QCM-D and AFM findings, attention to the differences between mica and silica must be paid. Mica is a crystalline aluminosilicate, while the silica used in QCM-D is amorphous silicate. Additionally, differences in surface roughness exist; mica is nearly atomically smooth, while silica has height variations of ∼1 nm (potentially influencing CP adsorption). QCM-D interpretation is that no CP film was retained on silica; however, AFM and previously published SFA results^[6]^ of CP on mica in H_2_O show that CP indeed forms nanofilms on mica. Combined, these results refine our understanding of CP, a short hydrophobic peptide, interacting with substrates possessing strong hydration layers. In addition to the above-described differences between mica and silica, differences in sample preparation between QCM-D and AFM might also be considered. The silica QCM-D sensor continuously vibrates at MHz frequencies while CP is deposited, generating an oscillatory shear stress that may influence CP interaction with the hydrated silica surface, rather than the mica surfaces, which were incubated in a stationary state. The flow in QCM-D is laminar, fixed, unlike the “manual” deposition and rinsing for AFM samples, which might be an additional comparative consideration. Studying the impact of flow rate on CP film formation and retention might be an important future direction for optimizing the design of CP-functionalized surfaces for use as wet adhesives.

**Figure 6.**
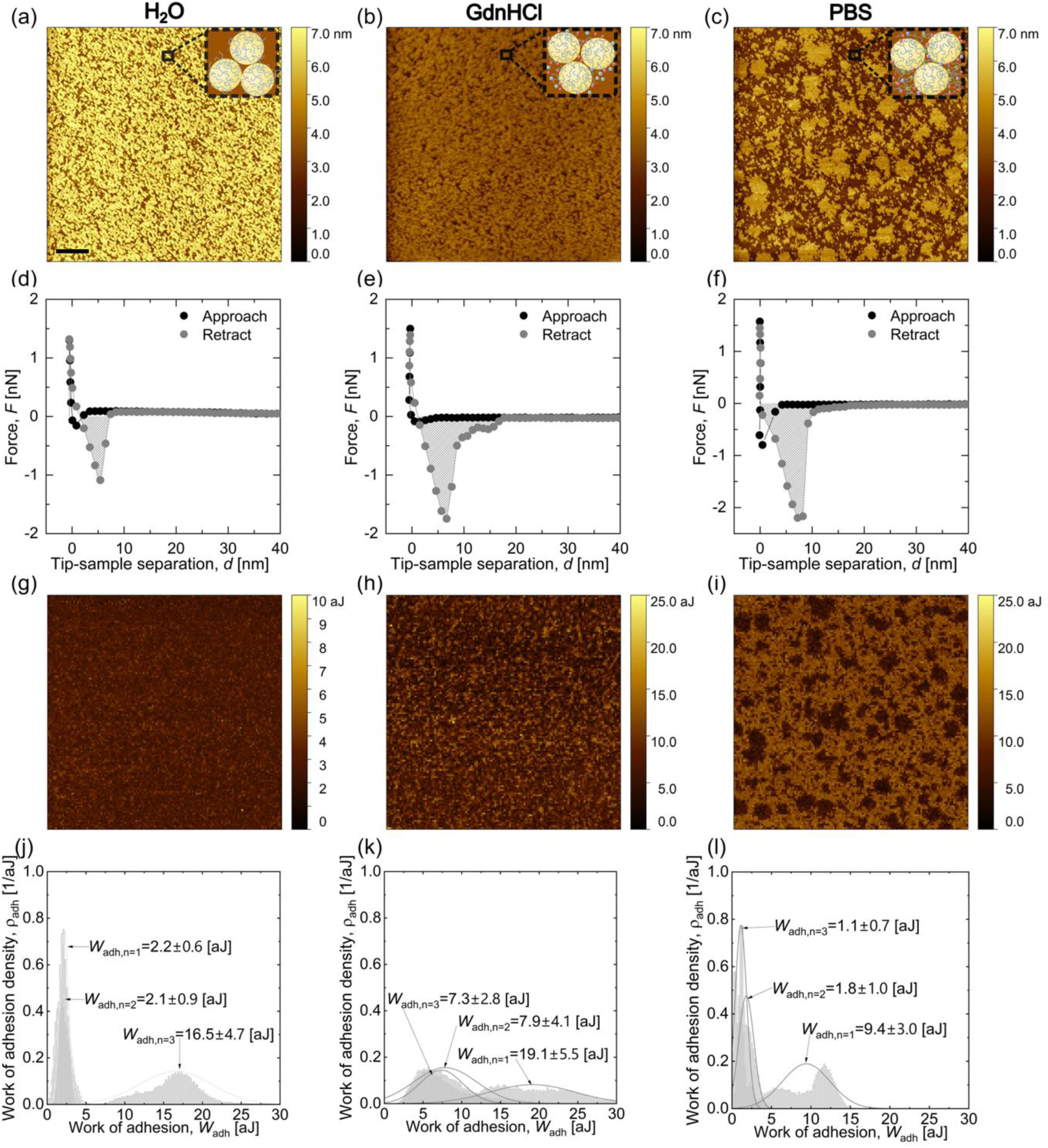
Representative AFM measurements of CP films on mica in different aqueous environments. Height maps of CP films, with schematic inserts illustrating potential molecular mechanism of CP aggregate deposition on mica, in (a) H_2_O, (b) GdnHCl, (c) and PBS, respectively. Representative force-runs in (d) H_2_O, (e) GdnHCl, (f) and PBS, respectively. Work of adhesion (*W*_adh_) maps, with optimized color scales for visualization of local differences, of CP films in (g) H_2_O, (h) GdnHCl (i) and PBS, respectively. (j), (k), (l) *W*_adh_ distribution histograms of CP films extracted from the *W*_adh_ from all measurements. Scale bar in (a) is 500 nm and applies to all AFM maps shown.

The representative FD curves for CP in H_2_O (Figure 6d) and in GdnHCl (Figure 6e) were similar, though the CP curve in GdnHCl showed a larger negative area. This is reflected in the *W*_adh_ maps of H_2_O (Figure 6g) and GdnHCl (Figure 6h), and *W*_adh_ histograms for H_2_O (Figure 6j) and GdnHCl (Figure 6k). For CP in PBS, in general, FD curves presented the following trend: out-runs of FD curves taken over AAMCs resulted in smaller *W*_adh_ compared to the rest of the surface. This is, higher *W*_adh_ values were quantified in regions between AAMCs (Figure 6l), potentially due to the presence of CP sub-nanofilms. In our previous work, we measured pull-off force between two CP films deposited on mica, in H_2_O and M9 (I∼190) buffer, with an SFA. M9 can be considered a kosmotrope, like the PBS used in this study, with effects on aggregation and strength of protein-based underwater adhesives.^[23]^ In those SFA experiments, for CP in M9, when approaching both surfaces into contact, two scenarios were encountered: *i*) a clean contact was possible, resulting in unambiguous FD curves and pull-off forces, and *ii*) presence of large aggregates, leading to double contacts with ambiguous FD curves and quantitatively unmeasurable pull-off forces. Combining this with our current findings, our interpretation for CP in PBS is as follows: a thin sub-layer of CP covered the mica surface, with AAMCs resting on the CP sub-layer. The smaller *W*_adh_ values measured over the AAMCs might be due to the convoluted effect of adhesion, weaker cohesion, and dissipative capabilities of the AAMCs, as they all contribute to *W*_adh_.

## CONCLUSIONS

To summarize, our work explored the impact of salts on the aggregation behavior of CP in aqueous solutions containing a cosolvent (DMSO) through DLS measurements. Micron-sized *D*_H_ measurements suggest that the formation of AAMCs was primarily driven by hydrophobic interactions between nonpolar moieties of CP, which may be trapped in DMSO-based molecular clusters. In line with this hypothesis, we investigated the film-forming behavior of these AAMCs by monitoring the frequency changes of CP-exposed crystal substrates with different surface chemistries (gold and silica) using QCM-D. Mass densities measured on CP-exposed gold and silica crystals indicated distinct film-forming characteristics. AAMCs preferentially adsorbed to gold crystals to form a rigid nanofilm, regardless of the type of salts present, resulting from hydrophobic collapse. However, AAMCs did not undergo hydrophobic collapse on silica surfaces. These results suggest that the film-forming behavior of AAMCs formed in bulk solutions is dominated by surface chemistry. Next, CP film morphologies and interfacial properties were measured by AFM in liquid environments upon their deposition on mica-templated gold and mica surfaces. On gold, AAMCs formed smooth nanofilms that closely matched the substrate’s surface morphology, with the condition measured in H_2_O yielding the highest *W*_adh_. The high *W*_adh_ of CP nanofilm in H_2_O suggests the existence of DMSO nanodomains formed by DMSO residues binding on nonpolar moieties on CP after hydrophobic collapse. In contrast, morphology results on mica substrate showed droplet-like CP aggregates in H_2_O and GdnHCl, suggesting that the integrity of CP aggregates was preserved during surface deposition. CP film morphology in PBS on mica shows features of both AAMCs and CP aggregates, suggesting bridging events between the cationic moiety of CP by multivalent anions. While the model substrates used in our study recapitulate a limited range of relevant engineering substrates, they enable mechanistic insights of CP-mediated film formation in different hydration environments on metallic (less hydrophilic) and silicate surfaces (more hydrophilic). Future research regarding the use of CP, a bacteria-derived short peptide containing multiple aromatic canonical amino acid moieties, or peptides with similar properties, in different wet adhesion applications should consider effects of solvent conditions and surface chemistry that could alter film formation behaviors, and therefore further affect their adhesive performance. More importantly, the resulting insights may inform the development principles of peptide-based adhesives and coatings for biomedical devices, marine technologies, or other materials operating in aqueous environments.

## EXPERIMENTAL SECTION

### Materials

DMSO (LC-MS grade, REF 85190) and 1X PBS (Gibco, REF 10010-023) were purchased from Thermo Fisher Scientific (USA) and used as received. Guanidine hydrochloride (GdnHCl) salts were purchased from Fisher Scientific (>= 99.5%, BP178-500, Fisher Bioreagents^TM^, USA). Syringe filters with a pore size of 0.2 µm for solvent prefiltration were purchased from Millipore Sigma (Millex^TM^ SLLGC25NS, USA). Crystal sensors (silica-coated and gold-coated) used for QCM-D were purchased from Quartz Pro. Two-part epoxy glues (Gorilla Glue), containing a bisphenol A-epichlorohydrin polymer (resin) and a hardener, were used to immobilize substrates for AFM measurements. Natural muscovite mica (V1 quality) was purchased from S&J Trading Inc. High-purity gold pellets (99.99%) purchased from Kurt J. Lesker Co. were used to thermally deposit gold on mica substrates. Disposable petri dishes (35 mm) were purchased from CELLTREAT Scientific Products.

### Peptide Synthesis

We synthesized the CP peptide, SYWFFGWHTK, in-house using the Standard Fmoc Solid-Phase Peptide Synthesis (SPPS) method on a CEM Liberty Blue microwave-assisted automated synthesizer. Full details of the peptide synthesis protocol, along with peptide molecular confirmation by Matrix-Assisted Laser Desorption Ionization Time-of-Flight (MALDI-TOF) mass spectroscopy (Bruker Microflex LRF) and high-pressure liquid chromatography (HPLC, Agilent Infinity II 1260 analytical), can be found in the supplemental information of our previous work.^[6]^ The purity of CP peptide from our in-house synthesis was at least 95% or higher.

### CP solution preparation

Ultrapure water was collected from a Thermo Fisher Scientific purification system with a resistivity of 18.2 MΩ. The pH of the water was adjusted to neutral using ∼1 M HCl or NaOH before adding GdnHCl salts to prepare a buffer solution (165 mM). A stock solution of CP was prepared by dissolving lyophilized CP in DMSO to a concentration of 150 M, yielding a transparent, homogeneous solution, and stored at -20 °C until used. Shortly before each experiment, a stock CP aliquot was thawed and diluted in H_2_O, GdnHCl or 1X PBS (160 mM) to 1.5 μM, the working CP molarity used for all experiments in this study. All solvents used for sample preparation (*i.e.*, DMSO, H_2_O, GdnHCl, and PBS) were pre-filtered using 0.2 μm hydrophilic syringe filter.

### Dynamic Light Scattering (DLS)

To measure the hydrodynamic diameter (*D*_H_) of the CP in different solvents, we used a Zetasizer Pro-Blue (Malvern Panalytical). The fluctuation in the backscattered light intensity from CP in different solutions was induced by a He-Ne laser at 633 nm and detected at 173 degrees. The Z-average *D*_H_ and polydispersity index (*PDI*) were then computed by the ZS XPLORER software from the autocorrelation function of the backscattered light intensity. Values reported were measured from at least three independent samples of each condition in disposable micro cuvettes (ZEN0040, Malvern Panalytical) at 25 *°*C. Statistical differences between conditions were determined by unpaired (two sample) *t*-test.

### Quartz Crystal Microbalance with Dissipation (QCM-D)

We performed QCM-D measurements using a Q-Sense Explorer microbalance (Biolin, Sweden). Sensors consisted of AT-cut quartz crystals with 5 MHz fundamental resonance frequency, coated with 50 nm of gold (Au) or silica (SiO2). All measurements were performed at 25 ^°^C, controlled by an incorporated Peltier unit. The outlet channel of the microfluidic chamber was connected to a peristaltic pump (ISMATEC), used to maintain a constant flow rate of 3.33 µL/sec.

The QCM-D experiment began with a fluid baseline (H_2_O, Guanidinium Chloride, or PBS) until a dispersion of 2 Hz/hr or less was achieved for each overtone. Next, 1.5 µM solutions of CP in the respective liquid solvent were flown by applying suction through the peristaltic pump until the fluidic chamber was saturated, and data were collected until a steady state was reached. This step is referred to as “adsorption” in the text. After this, a wash with the respective baseline fluid was performed, and the data collected from the steady state of this step is referred to as “retention”.

We studied film formation and desorption kinetics by fitting the changes in the frequency of each overtone, *Δf_n_*, with a Boltzmann (sigmoidal) function, implemented in Python:

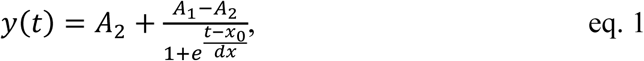

where *A*_2_ and *A*_1_ take the values of *Δf_n_* at the initial and final time of the film formation/desorption time domain, respectively. *x*_0_ is the inflection point of the fitted function (*i.e.*, where its second derivative ∼ 0), and *dx* a quantity that defines the transition steepness. We used nonlinear least-squares optimization with SciPy^[61]^ to determine the fitting parameters. *R*^2^ values of representative fits for each condition, along with their residuals, are shown in the SI document. After fitting, we obtained the film formation/desorption rates for each overtone by calculating the derivative of eq.1 with respect to time:

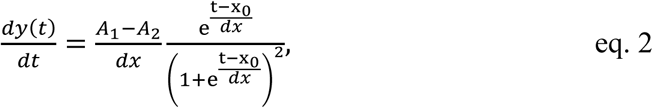

producing a Gaussian-like kinetic distribution that is centered at the inflection point *x*_0_. Next, we converted the rate curves (eq. 2) of each overtone into a normalized fractional completion parameter, mapping the formation/desorption processes into dimensionless progressions from 0 to 1. This is important, as it allows direct comparison in the formation/desorption times between different overtones, even when the absolute changes in frequency *Δf_n_* varies considerably among them. We did this by defining the function:

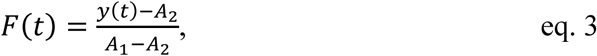

where *F*(0) = *A*_2_ and *F*(1) = *A*_1_, representing the initial and final states of film formation/desorption, respectively. Finally, we established film formation (Δ*t_f_*) / desorption (Δ*t_d_*) characteristic times as:

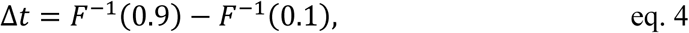

where *F*^−1^(*x*) is the inverse function of *F*(*x*), and thus *F*^−1^(0.9) and *F*^−1^(0.1) represent the times at which the film formation/desorption process reached 90% and 10%, respectively. We report the averages of Δ*t_f_* and Δ*t_d_* for overtones 3^rd^ to 13^th^ for each measurement. This procedure allows a non-biased determination of relative film formation/desorption times across measurements and minimizes potential measurement artifacts arising from the pump’s action at the first and last 10% of the formation/desorption process.

To calculate the average Sauerbrey mass (*m*_f_) of the adsorbed and retained films, we used the last 2 minutes of the steady state values (Δ*f*) for adsorption and retention. For all conditions, we report three independent measurements. We calculated *m*_f_ of the adsorbed films as the slope of normalized changes in frequency versus overtones, obtained by a linear regression from the 3^rd^ to the 13^th^ overtones^[62]^ following:

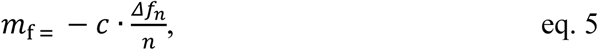

where *m*_f_ has units of ng/cm^2^, *c* is a constant that accounts for the crystal mass sensitivity (-17.7 ng·cm^-2^·Hz^-2^ for a 5 MHz crystal), and *Δf*_n_ is the change in frequency for the *n*-th overtone. For this purpose, we used the software pyQCM-BraTaDio.^[63]^ Statistical analysis of *m*_f_ was performed using two-way ANOVA in Origin(Pro) and unpaired (two sample) *t*-test.

### Atomic Force Microscopy (AFM)

For imaging and force mapping in liquid of the retained CP peptide films on mica and mica-templated gold substrates,^[64]^ we used a Nanowizard Pure AFM (Bruker, USA), equipped with ScanAsyst-Fluid probes, with a nominal spring constant of 0.7 N/m. We always calibrated each probe using the thermal-noise method^[65,66]^ prior to each measurement. We used QI imaging mode to obtain morphological maps (4×4 µm^2^). For force volume mapping, we used a setpoint of 2 nN, collecting a total of 16384 force-runs on each previously imaged area. For each condition with CP, at least 3 independent samples were probed.

Morphological images were processed using the software Gwyddion version 2.6.7.^[67]^ Row alignment (second-degree polynomial) and horizontal scar corrections were applied to the raw images. We report the work of adhesion (*W*_adh_), as the negative area of each out-run curve, using JPKSPM Data Processing version 8.1.62 and Origin(Pro) learning version.^[68]^ AFM measurements of bare substrates without CP in different solvents are displayed in supplemental information (Figures S10 to S14).

### CP film deposition for AFM experiments

We prepared two types of substrates for CP film deposition: mica and mica-templated gold. We immobilized freshly cleaved precut mica in a 35 mm disposable petri dish using a two-part epoxy glue. For the mica-templated gold substrate, we followed the previously published protocol^[64]^ using a benchtop thermal evaporator (Denton Vacuum, BTT IV). After thermal deposition of ∼ 50 nm of gold, we transferred the deposited gold layer to a silicon wafer substrate (∼ 1 x 1 cm^2^), leaving the smooth gold surface exposed. Next, we deposited CP on mica and mica-templated gold by drop-casting 50 -100 μL of CP solutions and allowed CP to adsorb for 30 minutes at room temperature. To minimize solvent evaporation during CP adsorption, we added droplets of the working solvent in the periphery, having the petri dish enclosed and parafilmmed. After incubation, we removed unbound CP from the surface and rinsed with the working solvent. For AFM measurements, we added 5 mL of the working solvent to the petri dish, allowing complete immersion of the CP films and AFM probes.

## ACKNOWLEDGMENTS

S.Z., A.D.M., and R.C.A.E. acknowledge funding from the National Science Foundation (NSF)-CREST: Center for Cellular and Biomolecular Machines through the support of the NSF Grant No. NSF-HRD-1547848. S.Z. acknowledges funding from the NIH G-RISE I-BioSTeP Grant No. T32 GM141862. R.C.A.E. and D.R.J.P. acknowledge funding from the Defense Advanced Research Projects Agency through the GLUE: Grip Likelihood in Underwater Environments Grant No. HR0011-24-3-03-62 awarded to R.C.A.E.. J.Y. acknowledges funding from the Defense Advanced Research Projects Agency through the GLUE: Grip Likelihood in Underwater Environments Grant No. HR0011-24-3-03-56. The views, opinions, and/or findings expressed are those of the authors and should not be interpreted as representing the official views or policies of the Department of Defense or the U.S. Government. J.Y. also acknowledges support from the Burroughs Wellcome Fund award No. 1022835. A.D.M acknowledges funding from the NSF DMR-2534247. T.P.C. acknowledges funding from the National Agency for Research and Development of Chile (ANID) through grant Fondecyt Regular 1251913. S.Z., D.R.J.P., and R.C.A.E. thank Dr. Mourad Sadqi for technical support and assistance during this project.

## SUPPLEMENTARY INFORMATION

**S1.** Estimation of CP monolayer mass density

**Table S1.** *D*_H_ of CP in DMSO, H_2_O, GdnHCl, and PBS.

**Table S2**. Unpaired *t*-test for *D*_H_ of CP in DMSO, H_2_O, GdnHCl, and PBS.

**Table S3.** PDI of CP in DMSO, H_2_O, GdnHCl, and PBS.

**Table S4**. Unpaired *t*-test for PDI of CP in different solutions.

**Figure S1.** DLS average hydrodynamic diameter of 1 vol. % DMSO in water.

**Figure S2.** *Δf* plots of CP on gold and silica.

**Figure S3.** *ΔD* plots of CP on gold and silica.

**Figure S4**. Boltzmann fit, transition rate, and residuals of fit of CP film formation on gold.

**Figure S5**. Boltzmann fit, transition rate, and residuals of fit of CP film formation on silica.

**Figure S6.** Boltzmann fit, transition rate, and residuals of fit of AAMC desorption on gold.

**Figure S7.** Boltzmann fit, transition rate, and residuals of fit of AAMC desorption on silica.

**Table S5.** Sauerbrey mass of CP adsorption and retention on gold and silica.

**Table S6.** Unpaired *t*-test analysis for Sauerbrey mass of CP adsorption and retention on gold and silica.

**Table S7.** Two-way ANOVA analysis for Sauerbrey mass of CP adsorption on gold and silica.

**Table S8.** Two-way ANOVA analysis for Sauerbrey mass of CP retention on gold and silica.

**Figure S8.** Decoupled step results of FD curves of CP on gold in H_2_O from mechanical mapping between CP-tip and DMSO-tip detachment.

**Figure S9.** AFM topography of mica-templated gold in H_2_O, 1 vol. % DMSO in H_2_O, 1 vol. % DMSO in GdnHCl, and 1 vol. % DMSO in PBS.

**Table S9.** Two-way ANOVA of RMS surface roughness of CP films on mica and mica-templated gold.

**Figure S10.** RMS roughness measurements of CP films on mica-templated gold and mica.

**Figure S11.** AFM topography of mica in H_2_O, 1 vol. % DMSO in H_2_O, 1 vol. % DMSO in GdnHCl, and 1 vol. % DMSO in PBS.

## Author contributions

The manuscript was written with contributions from all authors. All authors have approved the final version of the manuscript.

## REFERENCES

1. L. L. E. Mears, J. Appenroth, H. Yuan, et al., “Mussel Adhesion: A Fundamental Perspective on Factors Governing Strong Underwater Adhesion,” Biointerphases 17, no. 5 (2022), 10.1116/6.0002051.

2. C. Lee, H. Shi, J. Jung, et al., “Bioinspired Materials for Underwater Adhesion with Pathways to Switchability,” Cell Reports Physical Science 4, no. 10 (2023): 101597, 10.1016/j.xcrp.2023.101597.

3. H.-C. Flemming, J. Wingender, U. Szewzyk, P. Steinberg, S. A. Rice, and S. Kjelleberg, “Biofilms: An Emergent Form of Bacterial Life,” Nature Reviews Microbiology 14, no. 9 (2016): 563–575, 10.1038/nrmicro.2016.94.

4. L. Hall-Stoodley, J. W. Costerton, and P. Stoodley, “Bacterial Biofilms: From the Natural Environment to Infectious Diseases,” Nature Reviews Microbiology 2, no. 2 (2004): 95–108, 10.1038/nrmicro821.

5. J. K. Teschler, C. D. Nadell, K. Drescher, and F. H. Yildiz, “Mechanisms Underlying *Vibrio Cholerae* Biofilm Formation and Dispersion,” Annual Review of Microbiology 76, no. 1 (2022): 503–532, 10.1146/annurev-micro-111021-053553.

6. S. T. Ahmed, S. Zhai, X. Huang, et al., “Vibrio Cholerae Adhesin-Derived Peptides Mediate Strong Pull-off Forces in Aqueous Media with High Ionic Strength,” Colloids and Surfaces B: Biointerfaces 260 (2026): 115390, 10.1016/j.colsurfb.2025.115390.

7. X. Huang, T. Nero, R. Weerasekera, et al., “Vibrio Cholerae Biofilms Use Modular Adhesins with Glycan-Targeting and Nonspecific Surface Binding Domains for Colonization,” Nature Communications 14, no. 1 (2023): 2104, 10.1038/s41467-023-37660-0.

8. E. W. Danner, Y. Kan, M. U. Hammer, J. N. Israelachvili, and J. H. Waite, “Adhesion of Mussel Foot Protein Mefp-5 to Mica: An Underwater Superglue,” Biochemistry 51, no. 33 (2012): 6511–6518, 10.1021/bi3002538.

9. M. Varadi, D. Bertoni, P. Magana, et al., “AlphaFold Protein Structure Database in 2024: Providing Structure Coverage for over 214 Million Protein Sequences,” Nucleic Acids Research 52, no. D1 (2024): D368–D375, 10.1093/nar/gkad1011.

10. J. Jumper, R. Evans, A. Pritzel, et al., “Highly Accurate Protein Structure Prediction with AlphaFold,” Nature 596, no. 7873 (2021): 583–589, 10.1038/s41586-021-03819-2.

11. G. P. Maier, M. V. Rapp, J. H. Waite, J. N. Israelachvili, and A. Butler, “Adaptive Synergy between Catechol and Lysine Promotes Wet Adhesion by Surface Salt Displacement,” Science 349, no. 6248 (2015): 628–632, 10.1126/science.aab0556.

12. G. D. Degen, P. R. Stow, R. B. Lewis, et al., “Impact of Molecular Architecture and Adsorption Density on Adhesion of Mussel-Inspired Surface Primers with Catechol-Cation Synergy,” Journal of the American Chemical Society 141, no. 47 (2019): 18673–18681, 10.1021/jacs.9b04337.

13. H. Chang, V. Adibnia, W. Qi, R. Su, and X. Banquy, “Ternary Synergy of Lys, Dopa, and Phe Results in Strong Cohesion of Peptide Films,” ACS Applied Bio Materials 6, no. 2 (2023): 865–873, 10.1021/acsabm.2c01009.

14. H. Chang, V. Adibnia, C. Li, R. Su, W. Qi, and X. Banquy, “Short-Sequence Superadhesive Peptides with Topologically Enhanced Cation−π Interactions,” Chemistry of Materials 33, no. 13 (2021): 5168–5176, 10.1021/acs.chemmater.1c01171.

15. Y. Li, J. Cheng, P. Delparastan, et al., “Molecular Design Principles of Lysine-DOPA Wet Adhesion,” Nature Communications 11, no. 1 (2020): 3895, 10.1038/s41467-020-17597-4.

16. G. D. Degen, S. T. Ahmed, P. R. Stow, A. Butler, and R. C. Andresen Eguiluz, “PH-Tolerant Wet Adhesion of Catechol Analogs,” ACS Applied Materials and Interfaces 16, no. 17 (2024): 22689–22695, 10.1021/acsami.4c01740.

17. M. Shin, Y. Park, S. Jin, Y. M. Jung, and H. J. Cha, “Two Faces of Amine–Catechol Pair Synergy in Underwater Cation−π Interactions,” Chemistry of Materials 33, no. 9 (2021): 3196–3206, 10.1021/acs.chemmater.1c00079.

18. J. Gäding, V. Della Balda, J. Lan, et al., “The Role of the Water Contact Layer on Hydration and Transport at Solid/Liquid Interfaces,” Proceedings of the National Academy of Sciences 121, no. 38 (2024), 10.1073/pnas.2407877121.

19. J. Jacob, H. Duclohier, and D. S. Cafiso, “The Role of Proline and Glycine in Determining the Backbone Flexibility of a Channel-Forming Peptide,” Biophysical Journal 76, no. 3 (1999): 1367–1376, 10.1016/S0006-3495(99)77298-X.

20. P. Högel, A. Götz, F. Kuhne, et al., “Glycine Perturbs Local and Global Conformational Flexibility of a Transmembrane Helix,” Biochemistry 57, no. 8 (2018): 1326–1337, 10.1021/acs.biochem.7b01197.

21. M. Gruebele, K. Dave, and S. Sukenik, “Globular Protein Folding In Vitro and In Vivo,” Annual Review of Biophysics 45, no. 1 (2016): 233–251, 10.1146/annurev-biophys-062215-011236.

22. T. J. Gaborek, C. Chipot, and J. D. Madura, “Conformational Free-Energy Landscapes for a Peptide in Saline Environments,” Biophysical Journal 103, no. 12 (2012): 2513–2520, 10.1016/j.bpj.2012.11.001.

23. Z. D. Lamberty, C. M. Skogg, M. C. Wilson, et al., “Effect of Salts on the Aggregation and Strength of Protein-Based Underwater Adhesives,” ACS Omega 10, no. 45 (2025): 54535–54548, 10.1021/acsomega.5c07638.

24. R. M. Espinosa-Marzal, T. Drobek, T. Balmer, and M. P. Heuberger, “Hydrated-Ion Ordering in Electrical Double Layers,” Physical Chemistry Chemical Physics 14, no. 17 (2012): 6085, 10.1039/c2cp40255f.

25. R. M. Pashley, “DLVO and Hydration Forces between Mica Surfaces in Li+, Na+, K+, and Cs+ Electrolyte Solutions: A Correlation of Double-Layer and Hydration Forces with Surface Cation Exchange Properties,” Journal of Colloid and Interface Science 83, no. 2 (1981): 531–546, 10.1016/0021-9797(81)90348-9.

26. H.-N. Xu, Y. Liu, and L. Zhang, “Salting-out and Salting-in: Competitive Effects of Salt on the Aggregation Behavior of Soy Protein Particles and Their Emulsifying Properties,” Soft Matter 11, no. 29 (2015): 5926–5932, 10.1039/C5SM00954E.

27. A. M. Hyde, S. L. Zultanski, J. H. Waldman, Y.-L. Zhong, M. Shevlin, and F. Peng, “General Principles and Strategies for Salting-Out Informed by the Hofmeister Series,” Organic Process Research & Development 21, no. 9 (2017): 1355–1370, 10.1021/acs.oprd.7b00197.

28. M. C. Wilson, M. A. Beasley, K. P. Fears, E. A. Yates, and C. R. So, “Role of Protein Aggregate Structure on the Strength and Underwater Performance of Barnacle-Inspired Adhesives,” Soft Matter 19, no. 23 (2023): 4254–4264, 10.1039/D3SM00342F.

29. M. C. Wilson, Q. Lu, K. R. Nachtrieb, et al., “Underwater Adhesives Produced by Chemically Induced Protein Aggregation,” Advanced Functional Materials 34, no. 3 (2024), 10.1002/adfm.202308790.

30. S. Kim, H. Y. Yoo, J. Huang, et al., “Salt Triggers the Simple Coacervation of an Underwater Adhesive When Cations Meet Aromatic π Electrons in Seawater,” ACS Nano 11, no. 7 (2017): 6764–6772, 10.1021/acsnano.7b01370.

31. Q. Zhao, D. W. Lee, B. K. Ahn, et al., “Underwater Contact Adhesion and Microarchitecture in Polyelectrolyte Complexes Actuated by Solvent Exchange,” Nature Materials 15, no. 4 (2016): 407–412, 10.1038/nmat4539.

32. W. Wei, L. Petrone, Y. Tan, et al., “An Underwater Surface-Drying Peptide Inspired by a Mussel Adhesive Protein,” Advanced Functional Materials 26, no. 20 (2016): 3496–3507, 10.1002/adfm.201600210.

33. K. A. Ganar, M. Nandy, P. Turbina, et al., “Phase Separation and Ageing of Glycine-Rich Protein from Tick Adhesive,” Nature Chemistry 17, no. 2 (2025): 186–197, 10.1038/s41557-024-01686-8.

34. M. Karim, R. S. Boikess, R. A. Schwartz, and P. J. Cohen, “Dimethyl Sulfoxide (DMSO): A Solvent That May Solve Selected Cutaneous Clinical Challenges,” Archives of Dermatological Research 315, no. 6 (2022): 1465–1472, 10.1007/s00403-022-02494-1.

35. J. Stetefeld, S. A. McKenna, and T. R. Patel, “Dynamic Light Scattering: A Practical Guide and Applications in Biomedical Sciences,” Biophysical Reviews 8, no. 4 (2016): 409–427, 10.1007/s12551-016-0218-6.

36. V. Castelletto, I. W. Hamley, P. J. F. Harris, U. Olsson, and N. Spencer, “Influence of the Solvent on the Self-Assembly of a Modified Amyloid Beta Peptide Fragment. I. Morphological Investigation,” The Journal of Physical Chemistry B 113, no. 29 (2009): 9978–9987, 10.1021/jp902860a.

37. L. M. Carrick, A. Aggeli, N. Boden, J. Fisher, E. Ingham, and T. A. Waigh, “Effect of Ionic Strength on the Self-Assembly, Morphology and Gelation of PH Responsive β-Sheet Tape-Forming Peptides,” Tetrahedron 63, no. 31 (2007): 7457–7467, 10.1016/j.tet.2007.05.036.

38. Y. Feng, M. Taraban, and Y. B. Yu, “The Effect of Ionic Strength on the Mechanical, Structural and Transport Properties of Peptide Hydrogels,” Soft Matter 8, no. 46 (2012): 11723, 10.1039/c2sm26572a.

39. C. L. Shen, and R. M. Murphy, “Solvent Effects on Self-Assembly of Beta-Amyloid Peptide,” Biophysical Journal 69, no. 2 (1995): 640–651, 10.1016/S0006-3495(95)79940-4.

40. N. Habibi, N. Kamaly, A. Memic, and H. Shafiee, “Self-Assembled Peptide-Based Nanostructures: Smart Nanomaterials toward Targeted Drug Delivery,” Nano Today 11, no. 1 (2016): 41–60, 10.1016/j.nantod.2016.02.004.

41. K. L. Maxwell, D. Bona, C. Liu, C. H. Arrowsmith, and A. M. Edwards, “Refolding out of Guanidine Hydrochloride Is an Effective Approach for High-throughput Structural Studies of Small Proteins,” Protein Science 12, no. 9 (2003): 2073–2080, 10.1110/ps.0393503.

42. L. Platts, and R. J. Falconer, “Controlling Protein Stability: Mechanisms Revealed Using Formulations of Arginine, Glycine and Guanidinium HCl with Three Globular Proteins,” International Journal of Pharmaceutics 486, nos. 1–2 (2015): 131–135, 10.1016/j.ijpharm.2015.03.051.

43. C. S. Atwood, R. D. Moir, X. Huang, et al., “Dramatic Aggregation of Alzheimer Aβ by Cu(II) Is Induced by Conditions Representing Physiological Acidosis,” Journal of Biological Chemistry 273, no. 21 (1998): 12817–12826, 10.1074/jbc.273.21.12817.

44. Y. Bae, N. Nishiyama, S. Fukushima, H. Koyama, M. Yasuhiro, and K. Kataoka, “Preparation and Biological Characterization of Polymeric Micelle Drug Carriers with Intracellular PH-Triggered Drug Release Property: Tumor Permeability, Controlled Subcellular Drug Distribution, and Enhanced in Vivo Antitumor Efficacy,” Bioconjugate Chemistry 16, no. 1 (2005): 122–130, 10.1021/bc0498166.

45. H.-W. Sung, Y. Chang, I.-L. Liang, W.-H. Chang, and Y.-C. Chen, “Fixation of Biological Tissues with a Naturally Occurring Crosslinking Agent: Fixation Rate and Effects of PH, Temperature, and Initial Fixative Concentration,” Journal of Biomedical Materials Research 52, no. 1 (2000): 77–87, 10.1002/1097-4636(200010)52:1<77::AID-JBM10>3.0.CO;2-6.

46. P. Luo, Y. Zhai, E. Senses, et al., “Influence of Kosmotrope and Chaotrope Salts on Water Structural Relaxation,” The Journal of Physical Chemistry Letters 11, no. 21 (2020): 8970–8975, 10.1021/acs.jpclett.0c02619.

47. W. Koehnlein, E. Kastenmueller, T. Meier, T. Treu, and R. Falkenstein, “The Beneficial Impact of Kosmotropic Salts on the Resolution and Selectivity of Protein A Chromatography,” Journal of Chromatography A 1715 (2024): 464585, 10.1016/j.chroma.2023.464585.

48. A. M. S. Duarte, C. P. M. van Mierlo, and M. A. Hemminga, “Molecular Dynamics Study of the Solvation of an α-Helical Transmembrane Peptide by DMSO,” The Journal of Physical Chemistry B 112, no. 29 (2008): 8664–8671, 10.1021/jp076678j.

49. O. Zagorodko, T. Melnyk, O. Rogier, V. J. Nebot, and M. J. Vicent, “Higher-Order Interfiber Interactions in the Self-Assembly of Benzene-1,3,5-Tricarboxamide-Based Peptides in Water,” Polymer Chemistry 12, no. 23 (2021): 3478–3487, 10.1039/D1PY00304F.

50. H. Cui, M. J. Webber, and S. I. Stupp, “Self-assembly of Peptide Amphiphiles: From Molecules to Nanostructures to Biomaterials,” Peptide Science 94, no. 1 (2010): 1–18, 10.1002/bip.21328.

51. H. Fan, A. Rapikov, and E. Benassi, “Agglomeration in Dimethyl Sulfoxide and Water Binary System: A Comprehensive Study through Thermodynamics, Vibrational Spectroscopy, Quantum Mechanical Calculations and Morphology,” Journal of Molecular Liquids 408 (2024): 125333, 10.1016/j.molliq.2024.125333.

52. J. P. Gallivan, and D. A. Dougherty, “Cation-π Interactions in Structural Biology,” Proceedings of the National Academy of Sciences 96, no. 17 (1999): 9459–9464, 10.1073/pnas.96.17.9459.

53. Y. Akdogan, W. Wei, K. Huang, et al., “Intrinsic Surface-Drying Properties of Bioadhesive Proteins,” Angewandte Chemie International Edition 53, no. 42 (2014): 11253–11256, 10.1002/anie.201406858.

54. Y. Yang, J. Huang, D. Dornbusch, et al., “Effect of Surface Hydrophobicity on the Adsorption of a Pilus-Derived Adhesin-like Peptide,” Langmuir 38, no. 30 (2022): 9257–9265, 10.1021/acs.langmuir.2c01016.

55. S. Huang, Q. Hou, D. Guo, et al., “Adsorption Mechanism of Mussel-Derived Adhesive Proteins onto Various Self-Assembled Monolayers,” RSC Advances 7, no. 63 (2017): 39530– 39538, 10.1039/C7RA07425E.

56. W. Wei, L. Petrone, Y. Tan, et al., “An Underwater Surface-Drying Peptide Inspired by a Mussel Adhesive Protein,” Advanced Functional Materials 26, no. 20 (2016): 3496–3507, 10.1002/adfm.201600210.

57. N. R. Martinez Rodriguez, S. Das, Y. Kaufman, W. Wei, J. N. Israelachvili, and J. H. Waite, “Mussel Adhesive Protein Provides Cohesive Matrix for Collagen Type-1α,” Biomaterials 51 (2015): 51–57, 10.1016/j.biomaterials.2015.01.033.

58. I. Siretanu, S. R. van Lin, and F. Mugele, “Ion Adsorption and Hydration Forces: A Comparison of Crystalline Mica *vs.* Amorphous Silica Surfaces,” Faraday Discussions 246 (2023): 274–295, 10.1039/D3FD00049D.

59. K. Rechendorff, M. B. Hovgaard, M. Foss, V. P. Zhdanov, and F. Besenbacher, “Enhancement of Protein Adsorption Induced by Surface Roughness,” Langmuir 22, no. 26 (2006): 10885–10888, 10.1021/la0621923.

60. R. P. Richter, and A. Brisson, “QCM-D on Mica for Parallel QCM-DAFM Studies,” Langmuir 20, no. 11 (2004): 4609–4613, 10.1021/la049827n.

61. P. Virtanen, R. Gommers, T. E. Oliphant, et al., “SciPy 1.0: Fundamental Algorithms for Scientific Computing in Python,” Nature Methods 17, no. 3 (2020): 261–272, 10.1038/s41592-019-0686-2.

62. S. T. Ahmed, D. R. Jaramillo Pinto, L. Vitkova, et al., “Surface-Immobilized Fibronectin Conformation Influences Synovial Fluid Adsorption and Film Formation,” Colloids and Surfaces B: Biointerfaces 258 (2026): 115224, 10.1016/j.colsurfb.2025.115224.

63. B. Pardi, S. T. Ahmed, S. J. Flores, et al., “PyQCM-BraTaDio: A Tool for Visualization, Data Mining, and Modelling of Quartz Crystal Microbalance with Dissipation Data,” Journal of Open Source Software 9, no. 99 (2024): 6831, 10.21105/joss.06831.

64. L. Chai, and J. Klein, “Large Area, Molecularly Smooth (0.2 Nm Rms) Gold Films for Surface Forces and Other Studies,” Langmuir 23, no. 14 (2007): 7777–7783, 10.1021/la063738o.

65. J. L. Hutter, and J. Bechhoefer, “Calibration of Atomic-Force Microscope Tips,” Review of Scientific Instruments 64, no. 7 (1993): 1868–1873, 10.1063/1.1143970.

66. H.-J. Butt, and M. Jaschke, “Calculation of Thermal Noise in Atomic Force Microscopy,” Nanotechnology 6, no. 1 (1995): 1–7, 10.1088/0957-4484/6/1/001.

67. David Nečas, and Petr Klapetek, “Gwyddion: An Open-Source Software for SPM Data Analysis,” Cent. Eur. J. Phys. 10, no. 1 (2012): 181–188.

68. Origin(Pro), “Version 2026,” Version 2026, OriginLab Corporation: Northampton, MA, USA.

