## Supporting Information for "Surface-specific film assembly of a Vibrio cholerae adhesin-derived peptide modulated by environmental salts" (from another version of this preprint: version 1, 2026-06-23)



**S1.** Estimation of CP monolayer mass density
**Table S1.** $D_H$ of CP in DMSO, $H_2O$, GdnHCl, and PBS.
**Table S2**. Unpaired *t*-test for $D_H$ of CP in DMSO, $H_2O$, GdnHCl, and PBS.
**Table S3.** PDI of CP in DMSO, $H_2O$, GdnHCl, and PBS.
**Table S4**. Unpaired *t*-test for PDI of CP in different solutions.
**Figure S1.** DLS average hydrodynamic diameter of 1 vol. % DMSO in water.
**Figure S2.** $\Delta f$ plots of CP on gold and silica.
**Figure S3.** $\Delta D$ plots of CP on gold and silica.
**Figure S4**. Boltzmann fit, transition rate, and residuals of fit of CP film formation on gold.
**Figure S5**. Boltzmann fit, transition rate, and residuals of fit of CP film formation on silica.
**Figure S6.** Boltzmann fit, transition rate, and residuals of fit of AAMC desorption on gold.
**Figure S7.** Boltzmann fit, transition rate, and residuals of fit of AAMC desorption on silica.
**Table S5.** Sauerbrey mass of CP adsorption and retention on gold and silica.
**Table S6.** Unpaired *t*-test analysis for Sauerbrey mass of CP adsorption and retention on gold and silica.
**Table S7.** Two-way ANOVA analysis for Sauerbrey mass of CP adsorption on gold and silica.
**Table S8.** Two-way ANOVA analysis for Sauerbrey mass of CP retention on gold and silica.
**Figure S8.** Decoupled step results of FD curves of CP on gold in $H_2O$ from mechanical mapping between CP-tip and DMSO-tip detachment.
**Figure S9.** AFM topography of mica-templated gold in $H_2O$, 1 vol. % DMSO in $H_2O$, 1 vol. % DMSO in GdnHCl, and 1 vol. % DMSO in PBS.
**Table S9.** Two-way ANOVA of RMS surface roughness of CP films on mica and mica-templated gold.
**Figure S10.** RMS roughness measurements of CP films on mica-templated gold and mica.
**Figure S11.** AFM topography of mica in $H_2O$, 1 vol. % DMSO in $H_2O$, 1 vol. % DMSO in GdnHCl, and 1 vol. % DMSO in PBS.

**Table S1.** Summary table of averages and standard error of the means (SEM) for DLS $D_H$ of CP in DMSO, $H_2O$, GdnHCl, and PBS from three independent measurements (n).

| Solvent | Average $D_H$ | SEM | n |
|---|---|---|---|
| DMSO | 5 | 1 | 3 |
| $H_2O$ | 826 | 124 | 3 |
| GdnHCl | 1192 | 242 | 3 |
| PBS | 1167 | 94 | 3 |

**Table S2**. Summary table of unpaired *t*-test for DLS $D_H$ measurements of CP in DMSO, $H_2O$, GdnHCl, and PBS from three independent measurements (n).

| Group 1 | Group 2 | t-statistic | df | p-value | Result |
|---|---|---|---|---|---|
| $H_2O$ | PBS | -2.187 | 3.72 | 0.0991 | Not Significant ($p > 0.05$) |
| $H_2O$ | GdnHCl | -1.3433 | 2.98 | 0.2722 | Not Significant ($p > 0.05$) |
| $H_2O$ | DMSO | 6.6188 | 2 | 0.0221 | Significant ($p \leq 0.05$) |
| PBS | GdnHCl | -0.0981 | 2.59 | 0.929 | Not Significant ($p > 0.05$) |
| PBS | DMSO | 12.3911 | 2 | 0.0064 | Significant ($p \leq 0.05$) |
| GdnHCl | DMSO | 4.9003 | 2 | 0.0392 | Significant ($p \leq 0.05$) |

**Table S3.** Summary table of averages and standard error of the means (SEM) for DLS PDI of CP in DMSO, $H_2O$, GdnHCl, and PBS from three independent measurements (n).

| Solvent | Mean $D_H$ | SEM | n |
|---|---|---|---|
| DMSO | 0.48 | 0.1 | 3 |
| $H_2O$ | 0.56 | 0.1 | 3 |
| GdnHCl | 0.67 | 0.1 | 3 |
| PBS | 0.32 | 0.1 | 3 |

**Table S4**. Summary table of unpaired *t*-test for DLS PDI measurements of CP in different solutions from three independent measurements (n).

| Group 1 | Group 2 | t-statistic | df | p-value | Result |
|---|---|---|---|---|---|
| DMSO | $H_2O$ | -0.7648 | 3.35 | 0.4947 | Not Significant ($p > 0.05$) |
| DMSO | GdnHCl | -1.9946 | 3.42 | 0.1287 | Not Significant ($p > 0.05$) |
| DMSO | PBS | 2.3058 | 4 | 0.0824 | Not Significant ($p > 0.05$) |
| $H_2O$ | GdnHCl | -0.9986 | 3.99 | 0.3746 | Not Significant ($p > 0.05$) |
| GdnHCl | PBS | 3.764 | 3.42 | 0.0262 | Significant ($p \leq 0.05$) |
| $H_2O$ | PBS | 2.4868 | 3.34 | 0.0801 | Not Significant ($p > 0.05$) |

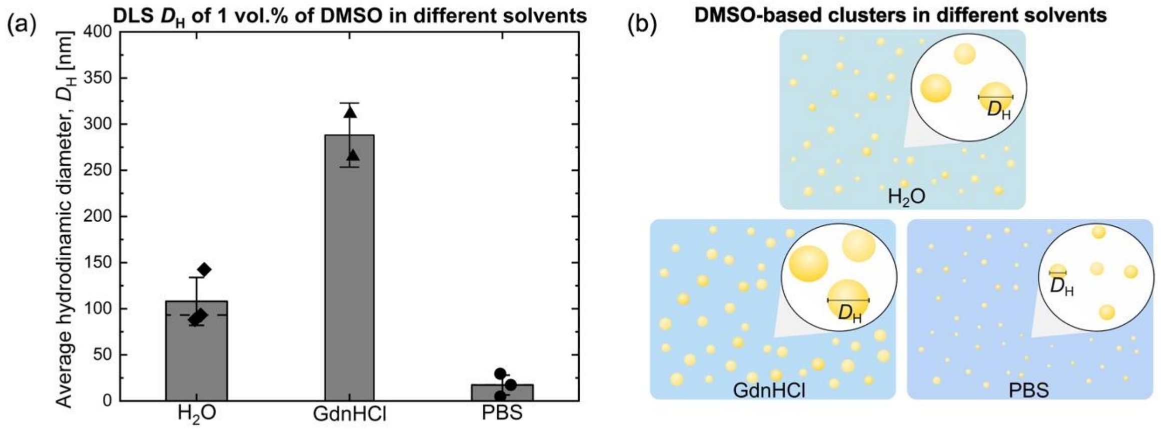


**Figure S1.** (a) DLS average hydrodynamic diameter measured in solvent mixtures containing 1 vol. % of DMSO. (b)Schematic diagram of DMSO-based clusters, with respective average $D_H$ in different solvent conditions, dispersed throughout the solvent.

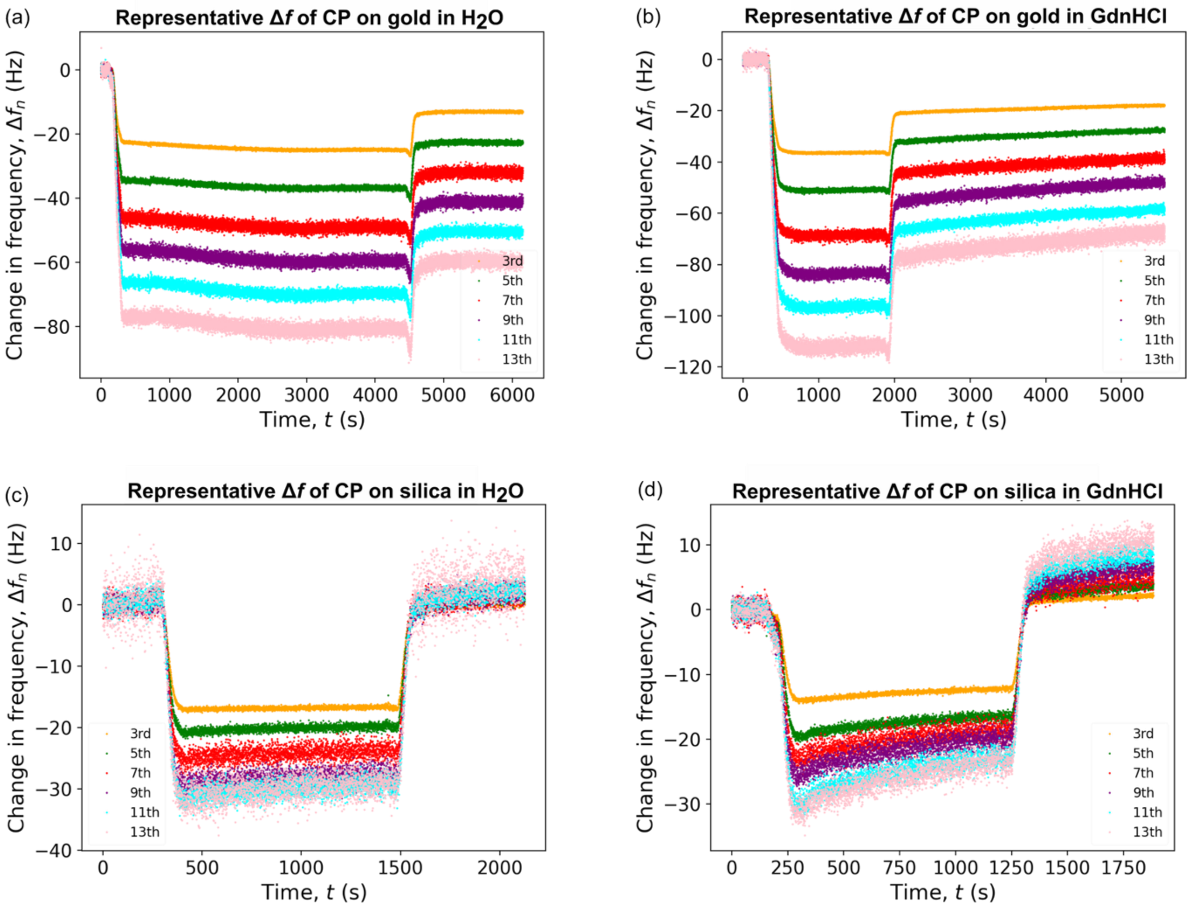


**Figure S2.** Representative *Δf* plots of CP on gold in $H_2O$ (a) and GdnHCl (b), showing CP films formed on gold substrates. In comparison, representative *ΔThe plots of CP on silica in H2O (c) and GdnHCl (d) showed the* absence of CP film formation.

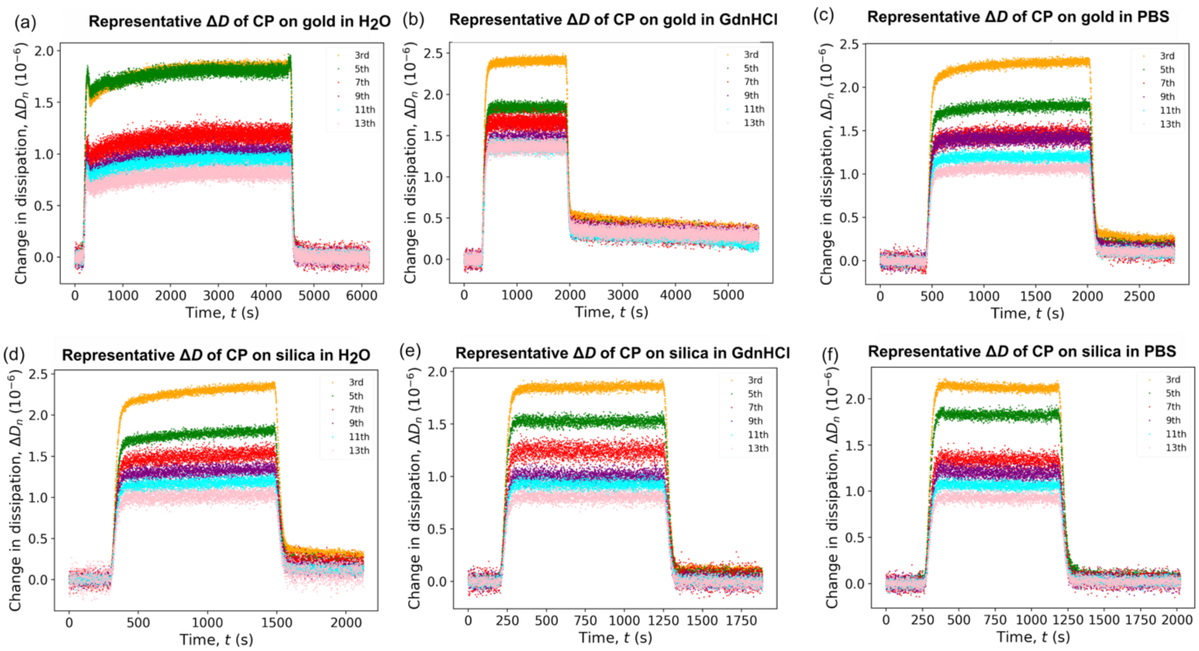


**Figure S3.** Representative *ΔD* plots of CP on gold in $H_2O$ (a), GdnHCl (b), and PBS (c) showing significant loss of dissipation on all overtones of CP film after wash. On the other hand, representative *ΔD* plots of CP on silica in $H_2O$ (d), GdnHCl (e), and PBS (f) also shown collapse of overtone dissipation, but without the formation of CP film.

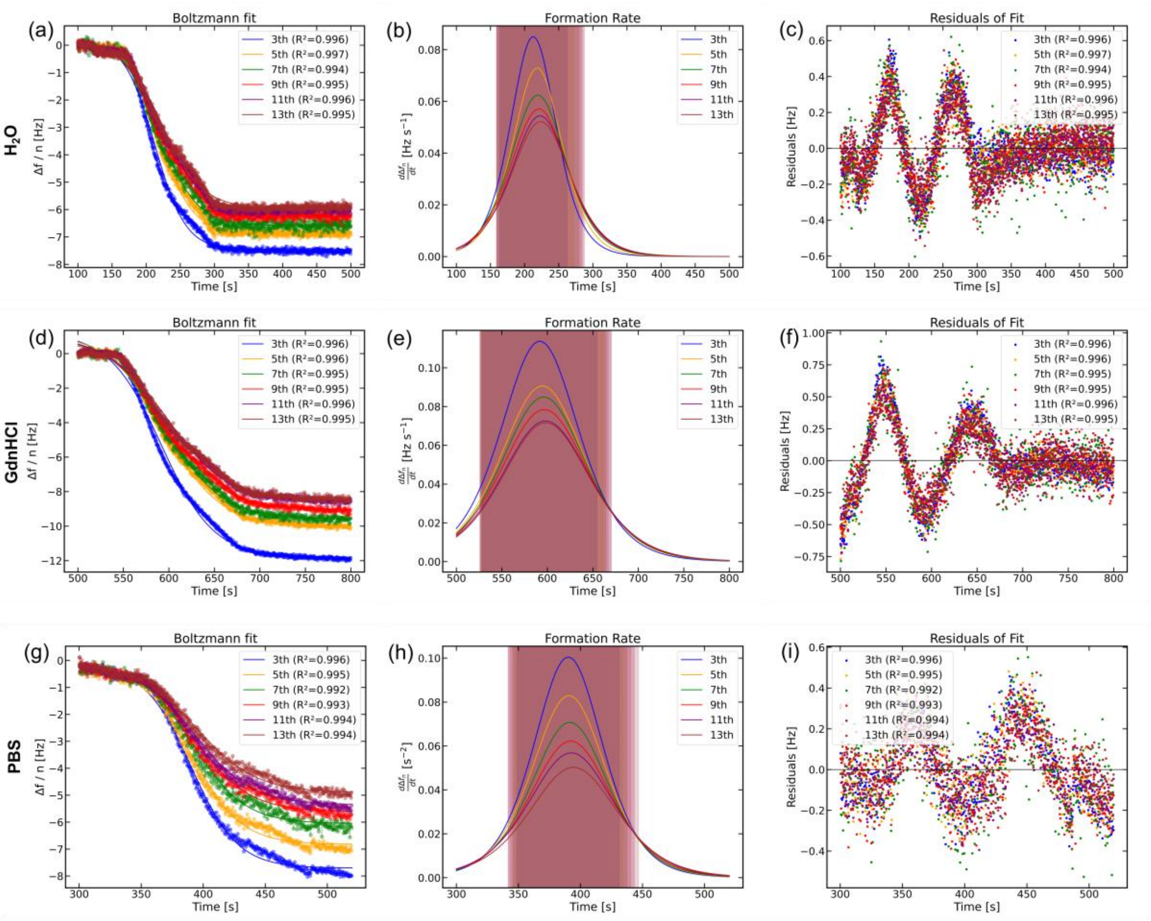


**Figure S4**. Representative Boltzmann fit, transition rate, and residuals of fit of CP film formation in $H_2O$ (a, b, & c), GdnHCl (d, e, & f), and PBS (g, h, & i), respectively, on gold.

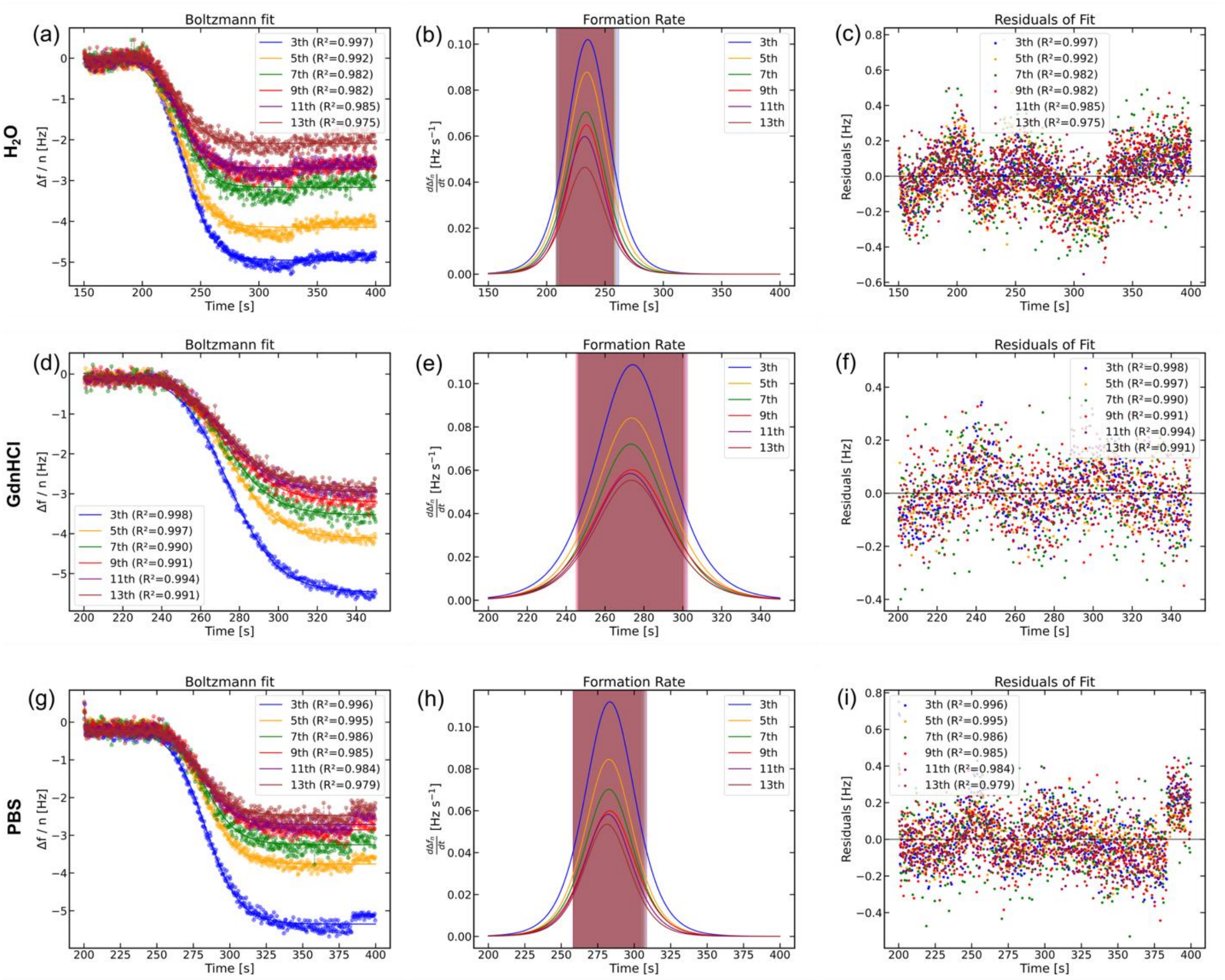


**Figure S5**. Representative Boltzmann fit, transition rate, and residuals of fit of CP film formation in $H_2O$ (a, b, & c), GdnHCl (d, e, & f), and PBS (g, h, & i), respectively, on silica.

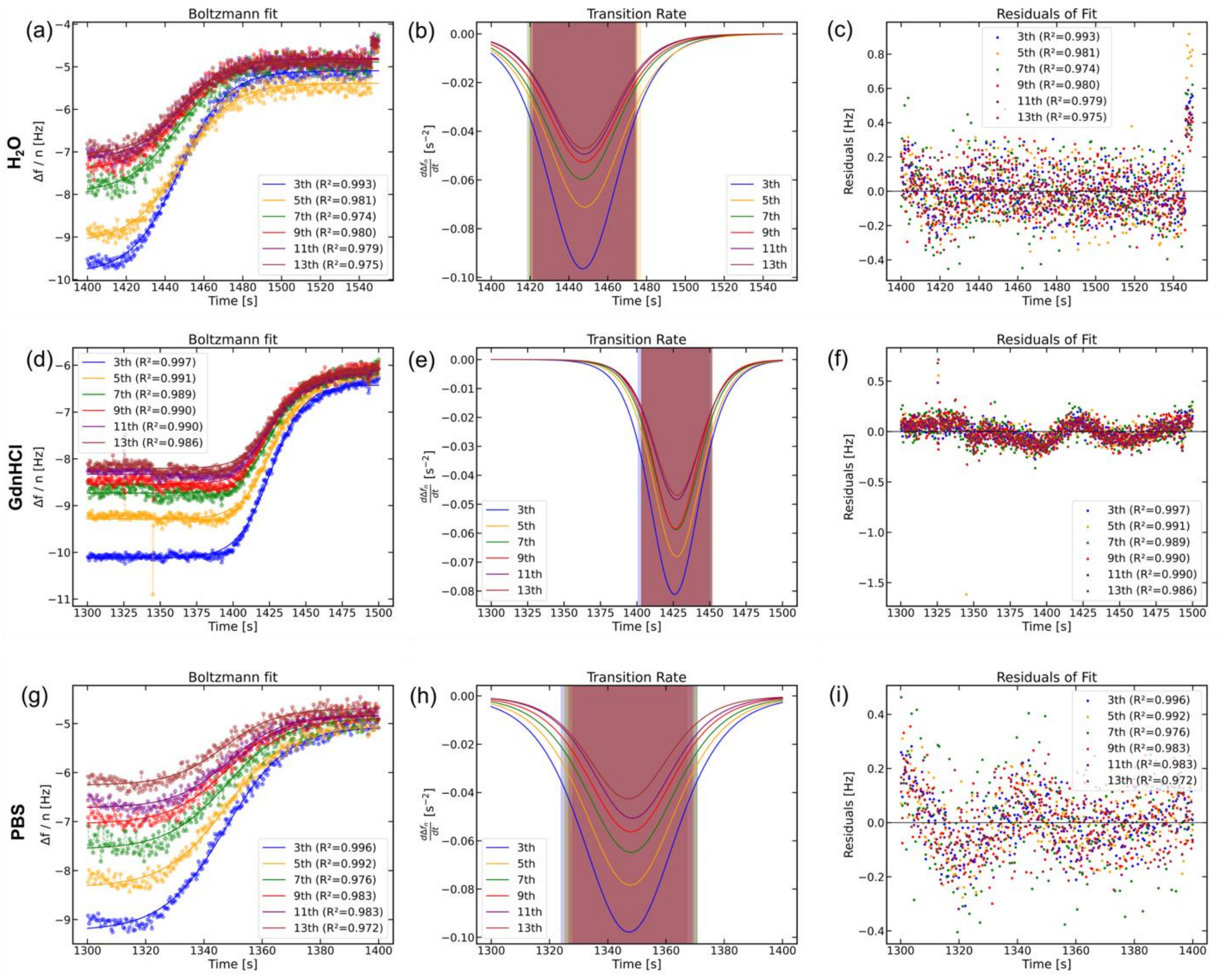


**Figure S6.** Representative Boltzmann fit, transition rate, and residuals of fit of AAMC desorption in $H_2O$ (a, b, & c), GdnHCl (d, e, & f), and PBS (g, h, & i), respectively, on gold.

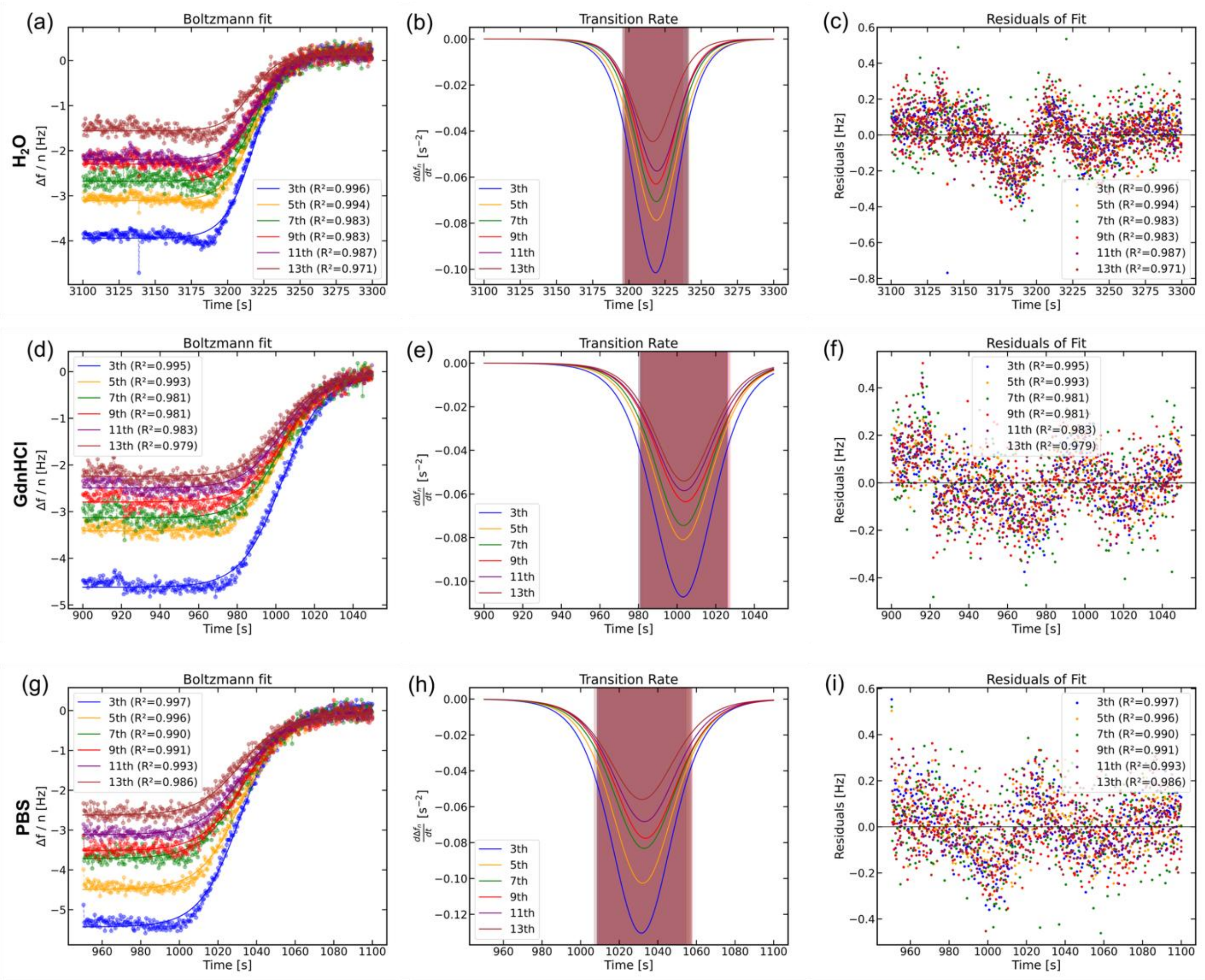


**Figure S7.** Representative Boltzmann fit, transition rate, and residuals of fit of AAMC desorption in $H_2O$ (a, b, & c), GdnHCl (d, e, & f), and PBS (g, h, & i), respectively, on silica.

**Table S5.** Summary table of averages and standard error of the means (SEM) for estimated Sauerbrey mass of CP adsorption and retention on gold- and silica-coated QCM crystals in DMSO, $H_2O$, GdnHCl, and PBS from three independent measurements (n).

| Substrate surface | Solvent | Mean | SEM | n |
|---|---|---|---|---|
| gold | $H_2O$ adsorption | 122.6 | 24.9 | 3 |
| gold | PBS adsorption | 100.0 | 14.2 | 3 |
| gold | GdnHCl adsorption | 140.8 | 7.6 | 3 |
| gold | $H_2O$ retention | 99.0 | 20.5 | 3 |
| gold | PBS retention | 92.8 | 11.2 | 3 |
| gold | GdnHCl retention | 77.6 | 18.8 | 3 |
| silica | $H_2O$ adsorption | 22.5 | 3.0 | 3 |
| silica | PBS adsorption | 27.3 | 1.9 | 3 |
| silica | GdnHCl adsorption | 25.2 | 2.9 | 3 |
| silica | $H_2O$ retention | 0.0 | 0.0 | 3 |
| silica | PBS retention | 0.9 | 0.9 | 3 |
| silica | GdnHCl retention | 0.0 | 0.0 | 3 |

**Table S6.** Summary table of unpaired *t*-test analysis for estimated Sauerbrey mass of CP adsorption and retention on gold- and silica-coated QCM crystals in DMSO, $H_2O$, GdnHCl, and PBS from three independent measurements (n).

| Group 1 | Group 2 | t-statistic | df | p-value | Result |
|---|---|---|---|---|---|
| $H_2O$ gold adsorption | $H_2O$ gold retention | 0.733 | 3.86 | 0.5056 | Not Significant ($p > 0.05$) |
| PBS gold adsorption | PBS gold retention | 0.3991 | 3.79 | 0.7113 | Not Significant ($p > 0.05$) |
| GdnHCl gold adsorption | GdnHCl gold retention | 3.121 | 2.64 | 0.0622 | Not Significant ($p > 0.05$) |
| $H_2O$ Silica adsorption | $H_2O$ Silica retention | 7.5833 | 2 | 0.0169 | Significant ($p \leq 0.05$) |
| PBS Silica adsorption | PBS Silica retention | 12.419 | 2.77 | 0.0017 | Significant ($p \leq 0.05$) |
| GdnHCl Silica adsorption | GdnHCl Silica retention | 8.6341 | 2 | 0.0132 | Significant ($p \leq 0.05$) |
| $H_2O$ gold adsorption | PBS gold adsorption | 0.7895 | 3.18 | 0.4844 | Not Significant ($p > 0.05$) |
| $H_2O$ gold adsorption | GdnHCl gold adsorption | -0.6982 | 2.37 | 0.5473 | Not Significant ($p > 0.05$) |
| PBS gold adsorption | GdnHCl gold adsorption | -2.5285 | 3.05 | 0.084 | Not Significant ($p > 0.05$) |
| $H_2O$ gold retention | PBS gold retention | 0.2667 | 3.1 | 0.8064 | Not Significant ($p > 0.05$) |
| $H_2O$ gold retention | GdnHCl gold retention | 0.7689 | 3.97 | 0.4851 | Not Significant ($p > 0.05$) |
| PBS gold retention | GdnHCl gold retention | 0.6924 | 3.27 | 0.5347 | Not Significant ($p > 0.05$) |
| $H_2O$ Silica adsorption | PBS Silica adsorption | -1.3444 | 3.45 | 0.2605 | Not Significant ($p > 0.05$) |
| $H_2O$ Silica adsorption | GdnHCl Silica adsorption | -0.6563 | 4 | 0.5475 | Not Significant ($p > 0.05$) |
| PBS Silica adsorption | GdnHCl Silica adsorption | 0.5796 | 3.48 | 0.5977 | Not Significant ($p > 0.05$) |
| $H_2O$ Silica retention | PBS Silica retention | -1 | 2 | 0.4226 | Not Significant ($p > 0.05$) |
| PBS Silica retention | GdnHCl Silica retention | 1 | 2 | 0.4226 | Not Significant ($p > 0.05$) |

**Table S7.** Summary table of two-way ANOVA analysis for estimated Sauerbrey mass of CP adsorption on gold- and silica-coated QCM crystals in DMSO, $H_2O$, GdnHCl, and PBS from three independent measurements (n).

| | DF | Sum of Squares | Mean Square | F Value | P Value |
|---|---|---|---|---|---|
| Solvent | 2 | 1129.44111 | 564.72056 | 1.15995 | 0.34188 |
| Substrate | 1 | 41596.89389 | 41596.89389 | 85.44112 | <0.0001 |
| Model | 3 | 42726.335 | 14242.11167 | 29.25367 | <0.0001 |
| Error | 14 | 6815.88111 | 486.84865 | | |
| Corrected Total | 17 | 49542.21611 | | | |

**At the 0.05 level, the population means of Solvent are not significantly different.**
**At the 0.05 level, the population means of Substrate are significantly different.**

**Table S8.** Summary table of two-way ANOVA analysis for estimated Sauerbrey mass of CP retention on gold- and silica-coated QCM crystals in DMSO, $H_2O$, GdnHCl, and PBS from three independent measurements (n).

| | DF | Sum of Squares | Mean Square | F Value | P Value |
|---|---|---|---|---|---|
| Solvent | 2 | 370.66778 | 185.33389 | 0.45143 | 0.64567 |
| Substrate | 1 | 36055.07556 | 36055.07556 | 87.82138 | <0.0001 |
| Model | 3 | 36425.74333 | 12141.91444 | 29.57475 | <0.0001 |
| Error | 14 | 5747.70111 | 410.55008 | | |
| Corrected Total | 17 | 42173.44444 | | | |

**At the 0.05 level, the population means of Solvent are not significantly different.**
**At the 0.05 level, the population means of Substrate are significantly different.**

### S1. Estimation of CP monolayer mass density

CP is a 10 amino acid sequence, SYWFFGWHTK. We can estimate the area of contact of one individual CP peptide, $A_{\mathrm{CP}}$, to be ~ 4 $\mathrm{nm}^2$. The molecular weight, $M_{\mathrm{w}}$, of CP is ~ 1.359 kDa, from where the mass density of a monolayer can be calculated as:

$$m_{monolayer} \approx \frac{M_{\mathrm{w}}}{A_{\mathrm{CP}}N_{\mathrm{A}}}\frac{[\mathrm{kDa}]}{\left[\frac{\mathrm{nm}^2}{\mathrm{mol}}\right]} = \frac{1359\left[\frac{\mathrm{g}}{\cancel{\mathrm{mol}}}\right]}{4\cdot 6.02\cdot 10^{23}\left[\frac{\mathrm{nm}^2}{\cancel{\mathrm{mol}}}\right]} \approx 56\left[\frac{\mathrm{ng}}{\mathrm{cm}^2}\right].$$

Doing the same calculation for an estimate $A_{\mathrm{CP}}$ ~ 3 $\mathrm{nm}^2$ gives $m_{monolayer}$ ~ $70\left[\frac{\mathrm{ng}}{\mathrm{cm}^2}\right]$, while $A_{\mathrm{CP}}$ ~ 5 $\mathrm{nm}^2$ gives $m_{monolayer}$ ~ $45\left[\frac{\mathrm{ng}}{\mathrm{cm}^2}\right]$. The $m_{\mathrm{f}}$ values in QCM-D for gold after the washing steps in all solvent system are close to these ranges, suggesting that CP retained on gold formed fully covering, dense monolayers.

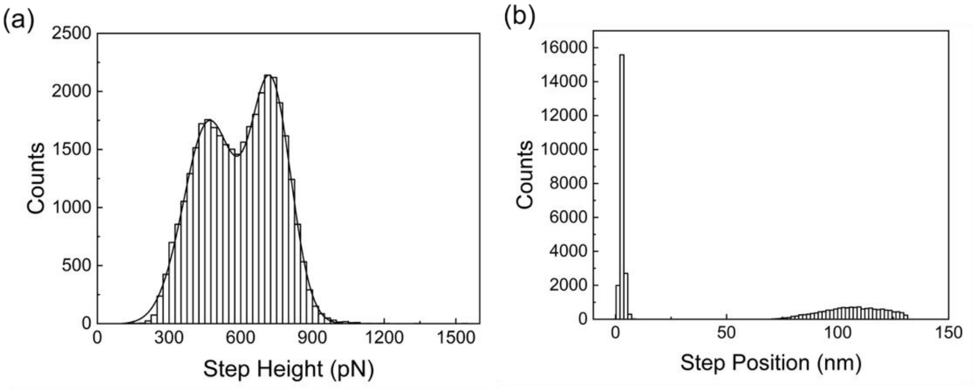


**Figure S8.** Decoupled step results of FD curves of CP on gold in $H_2O$ from mechanical mapping, showing two populations of jump-out forces (a) and positions (b) between CP-tip and DMSO-tip detachment; The adhesion forces of CP-tip and DMSO-tip detachment are different; one is around 400 pN and the other 700pN, respectively. The positions of the jump-outs are at 0.2 nm and 108 nm for CP-tip and DMSO-tip, respectively.

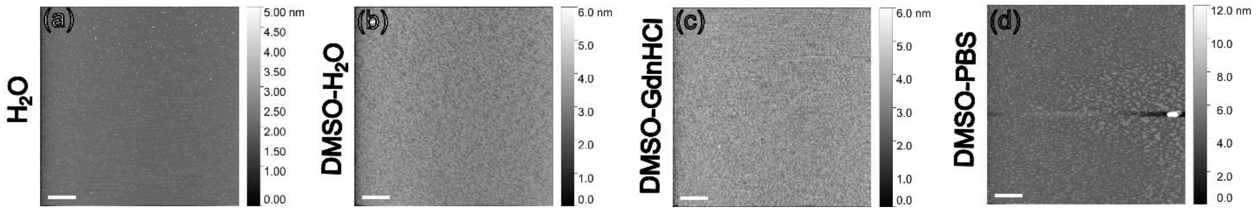


**Figure S9**. Representative AFM topography maps mica-templated gold in $H_2O$, 1 vol. % DMSO+$H_2O$, 1 vol. % DMSO+GdnHCl, and 1 vol. % DMSO+PBS environments. Scale bars are 500 nm.

**Table S9.** Summary table two-way ANOVA results comparing RMS surface roughness of CP films on mica and mica-templated gold in different solvents (e.g., $H_2O$, GdnHCl, and PBS), analyzed using Origin(Pro).

| | DF | Sum of Squares | Mean Square | F Value | P Value |
|---|---|---|---|---|---|
| Solvent | 2 | 0.14186 | 0.07093 | 0.55651 | 0.58391 |
| Substrate | 1 | 1.27633 | 1.27633 | 10.01374 | 0.00601 |
| Model | 3 | 1.46494 | 0.48831 | 3.83116 | 0.03046 |
| Error | 16 | 2.03933 | 0.12746 | | |
| Corrected Total | 19 | 3.50427 | | | |

At the 0.05 level, the population means of Solvent are not significantly different.
At the 0.05 level, the population means of Substrate are significantly different.

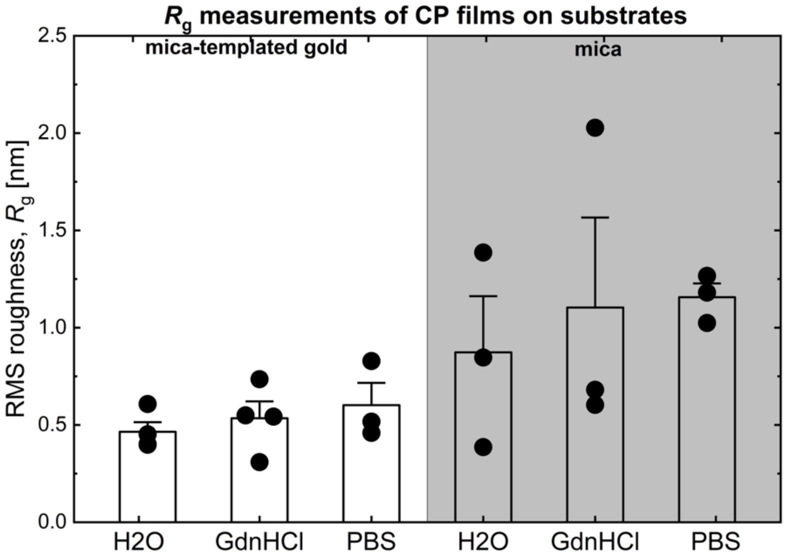


**Figure S10.** RMS roughness measurements of CP films on mica-templated gold and mica in different solvents (e.g., $H_2O$, GdnHCl, and PBS).

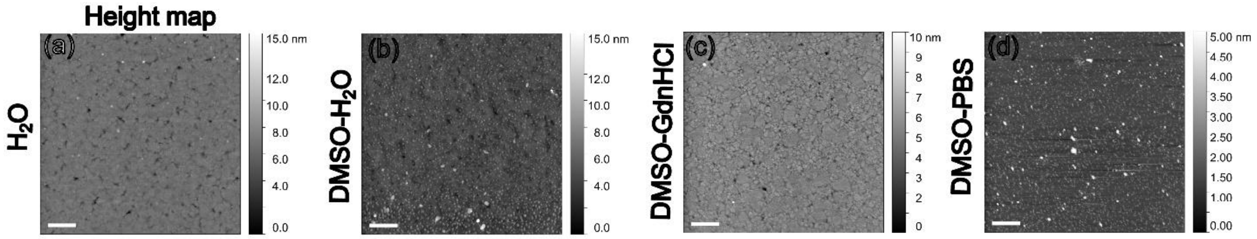


**Figure S11**. Representative AFM topography maps mica in $H_2O$, 1 vol. % DMSO+$H_2O$, 1 vol. % DMSO+GdnHCl, and 1 vol. % DMSO+PBS environments. Scale bars are 500 nm.
